# Spheroid surgery: a first-generation micromanipulation platform for extracting viable cell clusters from the outer layers of hydrogel-embedded spheroids

**DOI:** 10.64898/2026.10.22.760885

**Authors:** Viktor Pal, Miroslav Maiurov, Mariachiara Stellato, Nora Hapek, Maria Harmati, Filippo Piccinini, Akos Diosdi, Peter Horvath

## Abstract

Three-dimensional (3D) cell culture systems have gained attention as *in vitro* models capable of reproducing the tissue-like architecture, spatial organization, and physiological morphology of *in vivo* tumors. Their ability to mimic native cell-cell and cell-matrix interactions has positioned 3D cell cultures as powerful models for studying tumor biology and drug responses. However, methods for the localized extraction of viable cells remain limited, as existing approaches often disrupt the original 3D structure, limit access to defined regions, or compromise cell viability, restricting the biological insight that can be achieved.

We present a manual, micropipette-based manipulation system designed to extract viable multicellular clusters from the outer layers of compact 3D tumor spheroids, embedded in biocompatible hydrogels. The extracted clusters remained fully functional, demonstrating that the procedure is sufficiently gentle to remove small multicellular units.

This technology enables the localized extraction of viable multicellular clusters from 3D in vitro models without complete dissociation of the parent spheroid. The method opens new opportunities for longitudinal studies, spatially targeted sampling and controlled manipulation of living 3D cultures. This capability marks a step toward a new era of 3D model manipulation, where researchers can add, remove, or replace specific regions of living multicellular systems with precision.

## 1. Introduction

In recent years, three-dimensional (3D) cultures have gained a central role in oncological research, emerging as experimental tools that bridge the gap between traditional 2D *in vitro* models and the biological complexity of *in vivo* tumors^1,2^. Although 2D cultures represented the standard experimental framework for years, they impose physiological conditions different from real tumors^3^. Limitations include adhesion to rigid plastic surfaces that alters cell morphology, artificial oxygen and nutrient exchanges, less interaction between adjacent cells, and the lack of extracellular matrix hinder the possibilities for accurate comparison^4^. On the other hand, 3D systems allow cells to self-organise into complex spatial structures, establishing a more realistic cell-cell interactions and a microenvironment closer to real tumors. Physiological oxygen, nutrient, and pH gradients tend to emerge, generating a heterogeneous microenvironment inside the 3D structure^5^. This favours subpopulations with peripheral proliferative, quiescent intermediate, and central necrotic or quiescent regions, which are typical characteristics of *in vivo* solid tumors^6^. This regional differentiation influences drug penetration and distribution, making 3D models particularly relevant for pharmacological studies and preclinical screening^7^.

Among the various 3D cell culture models (e.g. spheroids, organoids, tumoroids, assembloids), multicellular spheroids are among the most widely adopted systems, owing to their ability to retain key features of *in vivo* tumors, including cellular heterogeneity, spatially organised regions of proliferation, quiescence and hypoxia, and physiological barriers to drug penetration^8^. Besides, spheroids allow a highly standardised and reproducible analysis, enabling consistent measurements for long-term and repeated imaging-based monitoring, including under prolonged drug treatment, reinforcing the role of 3D cultures as translational research tools^9,10^. However, the highly compact and dense structure of tumor spheroids poses significant challenges for obtaining detailed biological insight, particularly when attempting to access or manipulate specific spatial domains.

Cell isolation from *in vitro* 3D models and solid tissues is traditionally based on disaggregation procedures combining enzymatic treatments and mechanical solicitations to obtain a cell suspension suited for molecular and functional analysis^11^. The enzymatic digestion represents the most common approach and entails the use of proteases like trypsin, able to degrade the extracellular matrix components and the intercellular adhesion molecules^12,13^. This methodology has a high cellular yield, and it’s relatively easy to implement, but it requires highly optimised experimental parameters, such as long incubation times and high enzyme concentrations, which can hinder cellular viability, alter receptor integrity and induce transcriptional responses related to stress^14,15^. Mechanical dissociation, often used in combination with enzymatic methods, is based on the application of physical forces using repeated pipetting, trituration or filtration through a membrane with specific porosity^16–18^. Even though this method is faster and reduces enzymatic exposure, it also introduces transverse forces that can damage cells and affect the representativeness of isolated subpopulations. Another alternative is laser-based microdissection techniques that allow the selection of cells or cellular regions of interest directly in tissue sections through cutting or capture assisted by laser impulses^19–21^. This approach offers an elevated spatial precision, partially preserving the topological information of the culture^22^. Live-cell variants of laser microdissection have been described, but remain generally applied to fixed or cryopreserved samples, or to adherent monolayers, and are not readily suited to sampling defined regions of an intact 3D structure, limiting the possibility of functional analysis and re-culturing of material recovered directly from a spheroid^23^.

At the same time, the internal manipulation of 3D cell cultures^24^, such as being able to move cells in a controlled way, is increasingly recognised as a fundamental skill for future bioprinting and tissue engineering strategies^25^. Recent advancements in biofabrication technologies, like high-resolution bioprinting and micropatterning approaches, have focused on allowing the controlled repositioning of cells to recreate tissue architectures and cell-cell interaction networks^26,27^. This level of spatial control is fundamental for assembling tissues composed of multiple cell types, where relative positioning governs key processes for the culture’s microenvironment^28^. These types of technologies are expected to play a vital role in the generation of complex 3D organ-like tissues that can be used for different applications, ranging from advanced disease models and drug screening to the generation of implantable tissue constructs for regenerative medicine^29^.

Looking at the limitations of conventional extraction techniques, the necessity of developing new strategies capable of high spatial precision, morphological structure preservation, and cell viability maintenance arises. In this context, we propose a micropipette-based extraction system using pressure-controlled, borosilicate micropipettes, allowing the selection and extraction of multicellular clusters based on their position and morphological properties. This approach, which uses 3D spheroids embedded in biocompatible hydrogel for mechanical stability, allows controlled access to cells from the external layers while preserving the structural integrity of the parent spheroid and maintaining viability within the extracted cells. Unlike common bulk disaggregation techniques, it allows the possibility of performing multiple extractions from the same culture. Moreover, the integration with real-time imaging techniques paves the way to repeatable and reproducible sampling. It represents a first step to a minimally invasive and spatially resolved extraction protocol with future development aimed at single-cell resolution, access to deeper spheroid layers, and process automation.

## 2. Methods

### 2.1 Cell cultures

Two cell lines were chosen for the experiments, namely HeLa Kyoto expressing EGFP-α-tubulin and H2B-mCherry (Cell Lines Service, Eppelheim, Germany), a human cervical adenocarcinoma cell line, and T-47D (ATCC, Manassas, Virginia, United States), a human breast cancer cell line. HeLa Kyoto cells were maintained following the guidelines provided by the distributor, using cell culture medium consisting of DMEM (Corning, New York, United States), supplemented with 10% Fetal Bovine Serum (FBS, Corning, New York, United States), 2 mM L-glutamine (Gibco, Montana, United States), 0.5 mg/ml G418 (Gibco, Montana, United States), and 0.5 µg/ml Puromycin (Sigma-Aldrich, Missouri, United States). For T-47D, RPMI medium (Corning, New York, United States), supplemented with 10% FBS, 1% Penicillin-Streptomycin-Amphotericin B (Gibco, Montana, United States) and insulin (Gibco, Montana, United States) diluted to a final concentration of 0.2 units/ml, was used according to the manufacturer’s guidelines. Both cultures were maintained at 37 °C in a humidified incubator with 5% CO□.

For monoculture spheroid generation, the liquid overlay technique was applied to both cell lines using a hydrophobic surface-treated petri dish, namely the Cell Floater Dish (SPL Life Sciences Co., Ltd., Pocheon, South Korea). A new batch of spheroids was generated every three to four days by seeding approximately 800,000 cells per petri dish. Using this method, cell attachment occurred without spatial constraints, resulting in a heterogeneous and uneven cell distribution within the spheroids. Following seeding, the spheroids were used for experiments between days 1 and 4. During this period, they were maintained at 37 °C in a humidified incubator with 5% CO□.

To assess the suitability of scaffold-free and scaffold-based configurations for cell-cluster extraction, spheroids generated in ultra-low-attachment conditions were either embedded in a hydrogel matrix or transferred onto untreated culture surfaces and allowed to attach partially for 1-h. This temporary immobilization provided a stable reference point for controlled micropipette-based extraction, while allowing both the parental spheroid and the extracted cell clusters to be recovered from the culture medium for subsequent analyses.

Spheroids were embedded in a synthetic hydrogel matrix using the TrueGel3D hydrogel system (Sigma-Aldrich, USA) under sterile conditions, with all procedures performed in a laminar flow hood. For the experiment, 1 to 4 days old spheroids were collected from a hydrophobic surface-treated petri dish using a standard manual pipette and transferred into sterile Eppendorf tubes.

### 2.2 Optimization of the hydrogel formulation

The hydrogel formulation was optimized in-house, starting from the manufacturer’s protocol (TrueGel3D, Sigma-Aldrich). The final mixture was prepared by sequentially combining the following components in a microcentrifuge tube: 8.55 µL Milli-Q water, 1.2 µL TrueGel3D buffer, 0.9 µL SLO–Dextran, 3 µL spheroid suspension, and 1.35 µL CD cell-degradable crosslinker. In accordance with the manufacturer’s instructions, the SLO–Dextran-to-crosslinker volume ratio (1:1.5) was kept constant to maintain the stoichiometric balance between the reactive groups of the polymer and the crosslinker. Each component was thoroughly mixed before the addition of the next to ensure a homogeneous hydrogel solution. The mixture was maintained at room temperature for 2–10 min, until a slight increase in viscosity was observed. Increasing the crosslinker volume above 1.35 µL (with SLO–Dextran adjusted proportionally) made the hydrogel more difficult to work with, although successful extractions were still achieved in some cases. The manufacturer’s protocol identifies the crosslinker as oxygen-sensitive; therefore, it was aliquoted to minimize repeated air exposure. Despite these precautions, batch-to-batch variability persisted.

Subsequently, 15 µL of the spheroid-containing hydrogel mixture was deposited into a sterile µ-Slide 1 Well Glass Bottom imaging chamber with a #1.5H glass coverslip (ibidi GmbH, Germany; Cat. No. 82107) to form a single hydrogel droplet. This chamber format is compatible with the Operetta high-content imaging system and was used for both micropipette-based manipulation and subsequent time-lapse imaging. The sample was incubated at 37 °C for 27.5 min to allow complete hydrogel polymerization. Following gelation, the hydrogel droplet was gently covered with a prewarmed culture medium, after which the sample was used for micropipette-based extraction and manipulation experiments.

### 2.3 Micro-pipette fabrication

Micropipettes were pulled from borosilicate glass capillaries using a dual-stage micropipette puller (PC-10, Narishige, Japan). Multiple combinations of heater settings were tested during method development; heater settings of 85.2 a.u. for the first pulling stage and 55.2 a.u. for the second stage were adopted because they consistently yielded tips within a functional range for the extraction procedure. Even with fixed puller settings, however, the resulting tip geometry varied between individual pulling cycles; this is an inherent characteristic of the puller itself rather than a deliberate per-sample adjustment, depending on the thermal states of the device and the environment. Each micropipette was therefore visually inspected under the microscope before use, and only pipettes with a suitable tip geometry were selected for the extraction experiments. The outer diameter at the micropipette tip was estimated retrospectively from recorded microscope videos using an empirically verified pixel scale and ranged from approximately 3.5 to 14 µm among the pipettes used in the experiments, with measurements remaining approximate because of image resolution and tip focus. Micropipettes were filled with Trypsin (0.25% Trypsin-EDTA, Sigma-Aldrich, USA; Cat. No. T4049) immediately before use.

### 2.4 Pressure-controller

Pressure at the micro-pipette tip was regulated using a custom-designed pressure controller based on an Arduino Uno microcontroller (Arduino AG, Italy), providing a pressure range from −500 mbar to +1000 mbar. Pressure parameters were likewise refined through preliminary trials prior to the experiments reported here. In the resulting protocol, positive pressure of approximately +1000 mbar was applied while the pipette tip was positioned above the sample during initial Z-approach; once the tip entered the hydrogel, pressure was reduced to the level of approximately +100 to +300 mbar to maintain outward flow of the trypsin filling the pipette and prevent clogging while advancing towards the target region. Increased pressure of +100-300 to +1000 mbar was applied only when visible clogging of the pipette tip was observed, or when pressure delivery appeared to be interrupted. As the pipette approached the target region, the positive pressure was reduced to approximately 0 mbar, and once the tip was in the immediate vicinity of the cell, negative pressure of approximately −300 to −500 mbar was applied to aspirate and extract cellular material. Pressure values were adjusted for each experiment and are reported for each evaluated extraction (**Supplementary Data 1**).

### 2.5 Microscopy and imaging setup

Micropipette-based extraction experiments were performed using a LNscope (Luigs & Neumann, Germany) equipped with a 40× water immersion objective (Olympus, Japan) and LN Junior Compact micromanipulator, Dodt gradient contrast imaging and a CMOS camera (ODC 825 microscope camera, KERN & SOHN GmbH, Germany).

Post-extraction imaging was performed on an Operetta CLS high-content imaging system (Revvity, formerly PerkinElmer, USA) using a 20× air objective (NA 0.4) with 2 × 2 binning (0.59 µm per pixel). Brightfield (5 ms exposure, 5% power) and DRAQ7 fluorescence (Alexa 647 channel; excitation 630 nm, emission 708 nm; 20 ms exposure, 50% power) images were acquired as z-stacks at a 4 µm step. The stack range was set with a margin below the parent spheroid so that extracted cells remained within the stack as they settled. Time-lapse imaging was subsequently performed to monitor cell movement, morphological changes, and cell division. Twenty-five time points were acquired at 1-h intervals, corresponding to a 24-h longitudinal observation period, under 37 °C and 5% CO□ conditions. For two extractions, time-lapse imaging was continued in a second measurement started within 1-h, for a further 20-h and 24-h, respectively (approximately 45-h and 48-h in total).

### 2.6 Extraction of multicellular clusters from the outer layers

Multicellular clusters were extracted from the outer layers of hydrogel-embedded HeLa Kyoto and T-47D spheroids using a combined enzymatic and mechanical procedure. A target region from any part of the outer layers of the spheroid was selected under real-time Dodt gradient-contrast imaging using the LNscope microscopy setup. Trypsin was locally delivered through the micropipette directly to the selected peripheral region of the spheroid to partially loosen cell-cell contacts before mechanical extraction. It should be noted that the relative contribution of enzymatic and hydrodynamic effects to this local loosening cannot be resolved with the present setup. The hydrogel is embedded in serum-containing medium, in which trypsin activity is expected to be rapidly inhibited, yet the enzyme is delivered within approximately ten micrometres of the target region, so a transient local activity before dilution and inhibition cannot be excluded. At the same time, the fluid flow emerging from the micropipette tip exerts a local shear force that may itself contribute to the disruption of cell–cell contacts.

Following local enzymatic treatment, the micropipette was brought into contact with the selected region, and controlled negative pressure was applied to detach cellular material from the spheroid surface. During extraction, typically only one or a few cells at the periphery of the detached material were partially aspirated into the micropipette tip, whereas the majority of the multicellular cluster remained outside the pipette lumen.

Micropipette tip diameter, pressure values, and the approximate size of the recovered cluster varied between experiments. For each evaluated extraction, the tip diameter, the pressure values and polarity of the approach, extraction and deposition steps, spheroid age, the number of recovered cells and, where available, the extraction time were recorded in a standardized log sheet (**Supplementary Data 1**); outcomes of all attempts are listed in **Supplementary Note 1**.

### 2.7 Manipulation of multicellular clusters

Following extraction, the recovered cluster was displaced away from the parent spheroid and released into the surrounding hydrogel by applying controlled positive pressure through the micropipette. The cluster was positioned at a sufficient distance from the parent spheroid to allow its unambiguous identification and longitudinal observation during subsequent imaging. This manipulation step was monitored and recorded in real time using the LNscope microscope equipped with Dodt gradient-contrast imaging, as described above. In separate experiments, the micromanipulator was repositioned and fixed at different orientations, allowing the micropipette to approach the spheroid from different directions; although repeated extractions from different regions of the same spheroid were not tested directly, this combination is technically feasible.

### 2.8 Post-extraction viability assays

Post-extraction viability was assessed based on plasma membrane integrity over the observation period. This criterion was chosen because the plasma membrane is directly exposed to mechanical load during aspiration, deposition, and release and therefore provides an immediate indicator of manipulation-induced damage. Active cell movement and, where detectable, cell division were recorded as additional indicators of retained viability and proliferative capacity. Following extraction and manipulation, samples were transferred to the Operetta high-content imaging system. Characteristic morphological features of the parent spheroid, visible in the brightfield channel, were used as spatial reference landmarks to relocate the extracted cluster within the surrounding hydrogel.

An initial assessment using Trypan blue delivered through the micropipette was not pursued further because the staining status of individual cells could not be reliably assessed, and the approach was unsuitable for longitudinal imaging. 20 µL of DRAQ7™ Dye (Invitrogen, Thermo Fisher Scientific, USA; Cat. No. D15105/D15106; 0.3 mM stock solution) was added to the 1980 µL of culture medium. The resulting final DRAQ7 concentration was 3 µM, calculated according to the total volume of culture medium in the imaging chamber. After the cluster was removed, the culture medium was aspirated from the well and 2 mL of dye-containing culture medium was carefully added to the sample.

The extracted cells and multicellular clusters were imaged using both brightfield and fluorescence channels. DRAQ7 is impermeable to cells with intact plasma membranes and produces a far-red nuclear fluorescence signal in membrane-compromised cells. Accordingly, cells exhibiting DRAQ7-associated fluorescence were classified as membrane-compromised, whereas cells remaining DRAQ7-negative were considered to have retained plasma membrane integrity at the corresponding observation time point^30^.

Brightfield images were used to assess cluster morphology, changes in cell position, and active cell movement, while the fluorescence channel was used to monitor DRAQ7 uptake. DRAQ7 uptake and proliferative behaviour were assessed for individually distinguishable cells within each recovered cluster whenever cell-level identification was possible. Cell movement and division were recorded as additional observations, with division providing evidence of retained proliferative capacity.

### 2.9 Image annotation and quantitative analysis

Post-extraction image series (25 time points at 1-h intervals over 24-h; 4 µm z-step) were manually annotated in napari v. 0.5.5^31^ by a single annotator using the Labels layer on the brightfield channel. At each time point, the in-focus z-plane of each object was selected based on boundary visibility, and its contour was delineated in that plane. Rounded cells with a discernible continuous outline were annotated individually, irrespective of DRAQ7 status; elongated objects with severely disrupted morphology and no continuous boundary were excluded. When individual cell boundaries could not be reliably distinguished, the entire cluster was annotated as a single object and analysed separately. For extractions imaged beyond 24-h, morphological measurements were restricted to the first 24-h to keep observation windows comparable across experiments.

Measurements were calculated from the labelled masks using a custom *Python* workflow based on *NumPy*, *pandas* and *scikit-image* (**Supplementary Software 1**), with spatial calibration taken from the Operetta acquisition metadata. Contour area and perimeter were measured in the selected focal plane, and circularity was calculated as 4πA/P². Net XY displacement was calculated as the Euclidean distance between the initial and final contour centroids. Signed Z focus displacement was calculated as the change in the absolute position of the selected focal plane, derived from acquisition metadata to account for changes in stack position; it therefore describes focal-plane displacement rather than reconstructed three-dimensional centroid movement. For experiments with several annotated cells, initial and final area and circularity, as well as Z focus displacement, were summarized as medians across cells, and XY displacement as a range.

The Operetta image datasets, corresponding annotation masks, and multichannel TIFF files used for the quantitative analysis are available in **Figshare** at https://doi.org/10.6084/m9.figshare.c.8785665.

### 2.10 Statistics and reproducibility

Extraction attempts and their outcomes were recorded individually in the standardized experiment log (**Supplementary Note 1**). An attempt was one manipulation directed at a target on a spheroid; a specimen was one spheroid-containing hydrogel sample. The longitudinal analysis comprised eight extraction experiments from seven specimens: six multicellular-cluster extractions and two sequential single-cell extractions from the same T-47D specimen. The latter were separate extraction events but shared one biological specimen. Results are descriptive. No hypothesis tests or inferential comparisons between cell lines were performed.

Extractions were classified by the intended target: an extraction aimed at a multicellular cluster was counted as a cluster extraction even if fewer cells were finally recovered. Each extraction is identified by specimen, type of the extracted material and attempt number (*YYYYMMDD_line_type_specimen_attempt*), consistent with **Supplementary Note 1**.

## 3. Results

In this work, we developed Spheroid surgery, a micromanipulation approach for displacing individual cells or small cell clusters from the periphery of tumor spheroids (Figure 1). Extraction is performed under direct optical control with a glass capillary that is advanced through the hydrogel and can reach the spheroid from multiple directions (Figure 1a,b). Such an approach allows to isolate targeted cells without disrupting the structure and overall viability, and follow the development of both the parent spheroid and the relocated cells (Figure 1c). Through careful parametrization of the model, the environment, and the extraction steps, singlets that separate from the extracted chain during displacement or compact clusters that stay attached to the contacted cell can be reliably extracted. As cells are removed mechanically, the integrity of the cell membrane can be monitored using DRAQ7 dye combined with long-term time-lapse imaging (Figure 1c).

**Figure 1.**
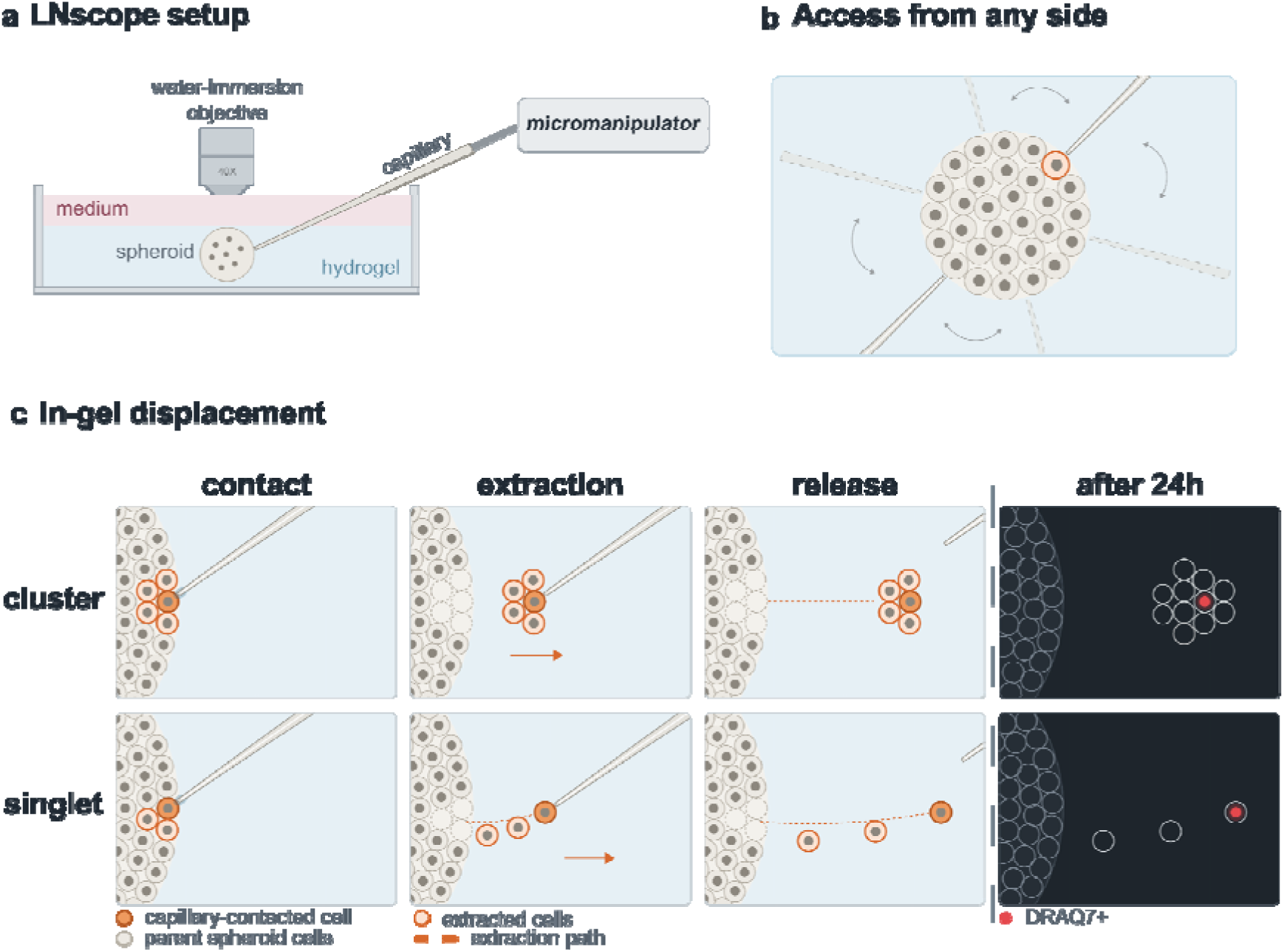
Schematic overview of spheroid surgery: in-gel displacement of single cells and cell clusters from hydrogel-embedded tumour spheroids. **a** Setup: the spheroid is embedded in TrueGel3D hydrogel under culture medium and imaged on an upright LNscope with a water-immersion objective (40×, NA 0.80); the glass capillary is mounted on a micromanipulator. **b** Approach directions around the spheroid. The glass capillary can reach cells from multiple directions, within the angle available between the sample holder and the objective. **c** Extraction sequence for a cluster (top row) and a singlet (bottom row). The first three columns show the brightfield steps: the capillary tip contacts a peripheral cell (contact), is moved laterally through the gel (extraction) and is withdrawn (release). The right column shows DRAQ7 fluorescence 24-h after extraction.

### 3.1 Impact of spheroid parameters on extraction reliability

Establishing reproducible cluster extraction required defining the structural properties of the spheroids and the embedding hydrogel that enable targeted detachment of cells without disrupting the parent structure. We therefore optimized the conditions (e.g. spheroid size, cell type and organization, maturation stage, and surface morphology) to standardize the micropipette-based extraction.

Spheroids below approximately 50–70 µm in diameter (typically <100 cells) were unsuitable for targeted removal of small clusters, as attempts to detach material frequently displaced the entire spheroid in our experimental system. Targeted cluster extraction was more feasible in spheroids containing more than approximately 100 cells and was consistently reproducible in spheroids exceeding 200 µm in diameter.

Extraction success was also influenced by cell type. Cell lines that form less dense and more loosely organized structure (e.g. HeLa Kyoto) were considerably easier to extract from. These spheroids typically exhibited an irregular surface morphology, with individual cells clearly distinguishable at the outer layers, facilitating micropipette access between cells and controlled detachment of cell clusters. In contrast, cell lines that form highly compact spheroids with smooth, continuous surfaces (e.g. T-47D, and MRC-5 in preliminary trials not included in the experiment log) required more careful manipulation during cluster extraction. Their dense architecture limited accessible entry points, making targeted extraction more challenging and often necessitating adjustments in enzyme delivery, pressure, and positioning.

Cluster extraction was also dependent on spheroid incubation time. As spheroids mature, they develop increasingly compact structures, making extraction progressively more difficult. Spheroids at earlier stages of incubation were generally easier to manipulate, possibly reflecting their less firmly established intercellular contacts. At 1 day post-seeding, spheroids had not yet developed sufficient structural integrity to support reliable cluster extraction, and removal attempts frequently resulted in disruption of the entire spheroid rather than selective detachment. A 2-4 day incubation window provided the optimal balance between structural stability and ease of targeted cluster extraction. During this period, spheroids remained sufficiently stable during manipulation, while retaining accessible regions where cell clusters could be reliably detached. Beyond this window, older spheroids exhibited highly compact cellular arrangements, which substantially reduced the likelihood of successful and controlled extraction.

Extraction conditions, including enzyme delivery, pressure, and targeting strategy, were adjusted according to cell line type, spheroid size, incubation time, and surface morphology. Detailed parameter adjustments and corresponding experimental outcomes are provided in the **Supplementary Note 1**.

### 3.2 External conditions affecting spheroid manipulation

Spheroids remained embedded in a hydrogel throughout extraction, and the properties of the surrounding matrix affected their manipulation. The matrix had to provide sufficient mechanical support for the spheroid while remaining penetrable by the micropipette. We therefore selected a synthetic hydrogel (TrueGel3D) to allow precise control of the physical properties. An additional advantage of this system is that the dextran-based backbone can be selectively degraded with a dedicated enzymatic solution (TrueGel3D Enzymatic Cell Recovery Solution, dextranase), allowing extracted cell clusters to be retrieved from the surrounding matrix for downstream analysis.

The hydrogel formulation was optimized (Section 2.2) by adjusting the amounts of dextran and crosslinker while maintaining their stoichiometric ratio and keeping the final hydrogel volume constant. This yielded a formulation that met both requirements. Hydrogels that were too stiff caused micropipette tip breakage and limited manoeuvrability, whereas formulations that were too soft failed to mechanically support the spheroid adequately and could also cause micropipette clogging, impairing pressure transmission. Following hydrogel optimization, successful cell-cluster extraction was reproducibly achievable in both HeLa Kyoto and T-47D cultures, although individual attempts could still fail because of target non-detachment, capillary clogging, or capillary breakage.

### 3.3 Demonstration of targeted cell-cluster extraction

The extraction procedure was evaluated based on the detachment and controlled release of cellular material, the outcomes of individual attempts, and the preservation of the parent spheroid structure. An attempt was considered successful when a single cell or a small cluster was detached from the spheroid surface and subsequently released at a distance from the spheroid that allowed its separate identification. Multiple attempts could be performed on the same specimen, and an experiment was classified as unsuccessful only if none met both criteria.

The outcomes of all extraction attempts and the final operating parameters are summarized in Figure 2. The experiment log documents 55 extraction attempts across 19 specimens, including method-development trials (Figure 2a). Twenty-six attempts were classified as technical failures (T), and 11 as extracted material that could not be evaluated further (C). Ten attempts resulted in successful extraction without subsequent dye screening (S), and eight were evaluated with DRAQ7 (E: six E+ and two E−; see Section 3.4 for the corresponding viability assessment). Because the log includes trials performed under changing conditions during protocol development, these pooled counts were not used to estimate a success rate for a fixed protocol. Instead, success was assessed in the final experimental series, performed once the protocol had been finalized (Figure 2c). In this series, 8 of 11 attempts yielded a single cell or cluster deposited in the hydrogel, and 5 of the 7 cells or clusters evaluated with DRAQ7 were classified as membrane-intact (E+). The three technical failures in this series were all capillary breakages that occurred in a single session (Figure 2b). All extraction attempts, including partially successful and unsuccessful ones and their specific technical limitations, are documented in **Supplementary Note 1** and the **Supplementary Videos**.

**Figure 2.**
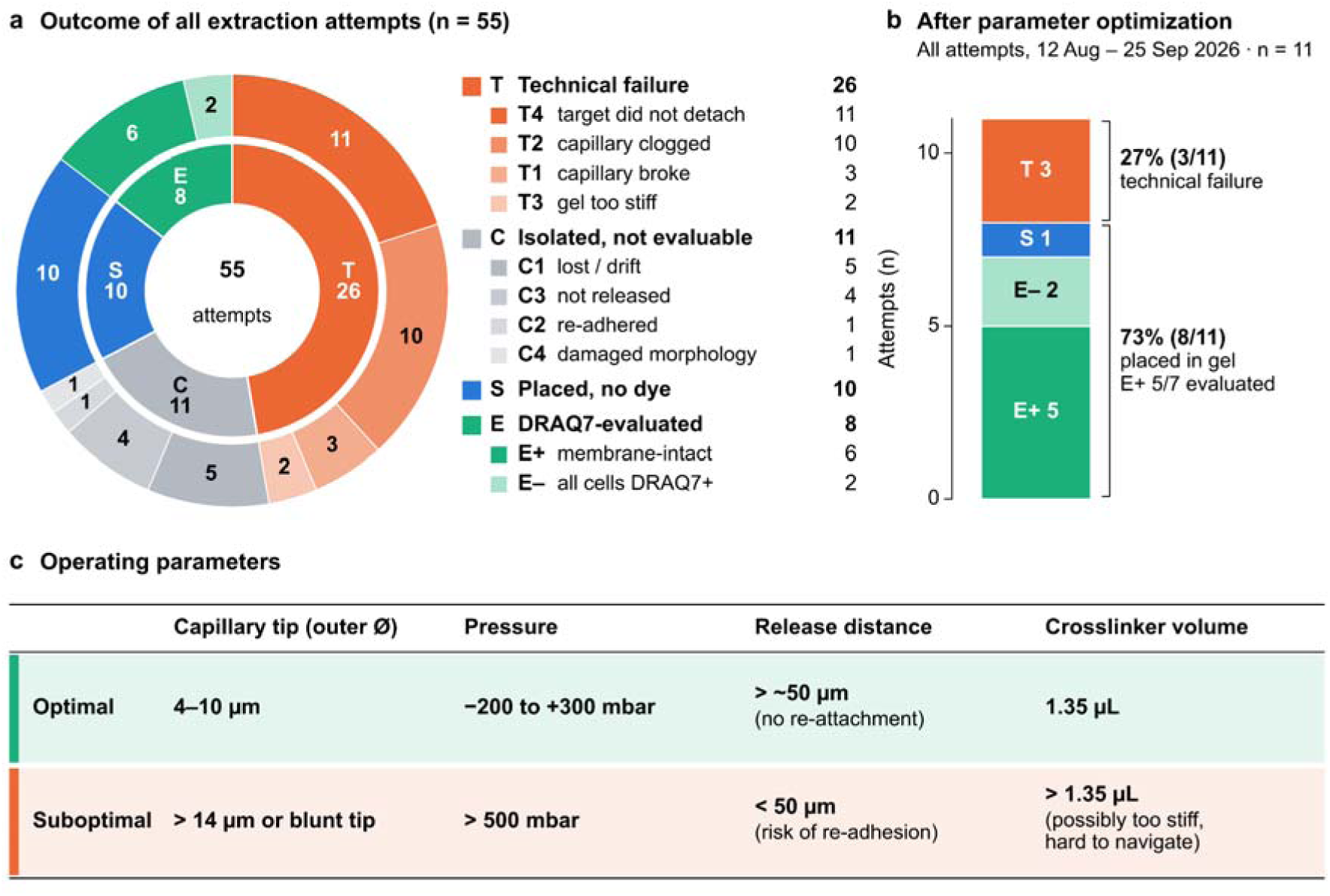
Extraction outcomes and optimized operating parameters. **a** Outcome of all extraction attempts (n = 55 attempts, 19 samples; HeLa Kyoto and T-47D). Inner ring, outcome class; outer ring, outcome code. T, technical failure; C, unit isolated but not evaluable; S, unit placed without viability dye; E, unit imaged with DRAQ7 for 23–24-h (E+, ≥1 cell DRAQ7-negative throughout; E–, all cells DRAQ7-positive). **b** Outcomes in the final series after parameter optimization (12 August – 25 September 2026; n = 11 attempts); E+ is given relative to DRAQ7-evaluated units. All technical failures were capillary breakages in one session. **c** Optimal and suboptimal operating parameters. Tip diameter, outer diameter measured against clear gel (±1 µm); release distance, gap between released unit and spheroid outline; crosslinker volume per 15 µL hydrogel.

Of the 18 successful extractions, eight, from seven specimens, were followed by DRAQ7 time-lapse imaging and evaluated in detail (E); the remaining ten were not imaged (S). The eight evaluated extractions comprised four HeLa Kyoto cluster extractions, two T-47D doublet extractions and two sequential single-cell extractions from one additional T-47D specimen. Cells were taken from the outer layers of the spheroid and deposited into the surrounding hydrogel (Figure 3). The extracted clusters initially contained approximately 2-15 cells. In every experiment, the parent spheroid retained its overall structure, with no visible structural disruption resulting from the extraction procedure. Fully detached clusters sometimes moved back towards the parent spheroid along the tunnel left by the capillary, usually within about 10 min; in the remaining cases, they stayed close to where they had been deposited.

**Figure 3.**
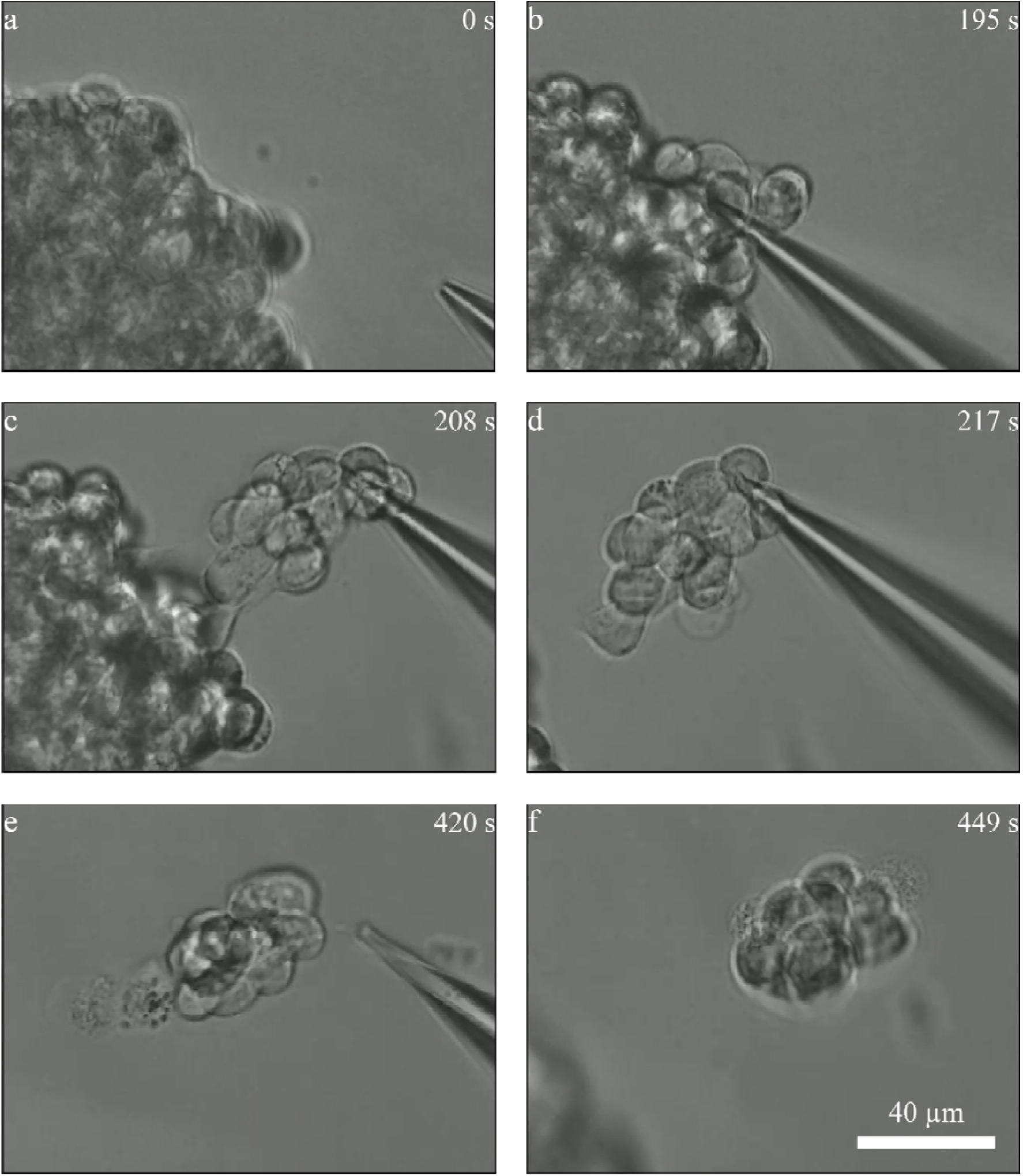
Micropipette-based extraction and manipulation of a multicellular cluster from the outer layer of a HeLa Kyoto spheroid embedded in hydrogel. (extraction *20260803_H_M_02_02*). a Micropipette positioning near the selected peripheral region of the spheroid. **b** Contact with the target region and initiation of aspiration. **c** Progressive detachment of the multicellular cluster from the spheroid surface. **d** Manipulation of the detached multicellular cluster using the micropipette. **e, f** Release of the extracted cluster from the micropipette. Each figure displays the time frame in seconds (s) in its top-right corner. The scale bar represents 40 µm.

Micropipette tip diameter varied between individual pipettes as an inherent consequence of the pulling process (Section 2.3). Pressure values were, in contrast, deliberately adjusted for each sample based on spheroid morphology, pipette tip size and hydrogel batch properties (**Supplementary Data 1**). Release distance was also adapted to individual samples (**Supplementary Note 1**). Consequently, the experiments were evaluated descriptively rather than as a formal parameter-optimization study.

### 3.4 Viability analysis of extracted multicellular clusters

Following extraction, membrane integrity was assessed using DRAQ7 staining, and the recovered cells were monitored by time-lapse imaging at 1-h intervals for at least 24-h. DRAQ7 imaging revealed heterogeneous membrane integrity within the extracted multicellular clusters: in several experiments, the cell directly contacted and aspirated by the micropipette became DRAQ7-positive, indicating membrane damage associated with the extraction procedure, whereas the remaining cells within the same cluster frequently remained DRAQ7-negative during the initial post-extraction observation period.

Cell division was observed in one of the six multicellular-cluster extraction experiments (Figure 4), with the first division detected approximately 6-h after the start of imaging. In two cluster extractions (*20260812_H_M_01_01* and *20260817_H_M_01_04*), time-lapse imaging was continued in a second measurement for a further 20-h and 24-h, respectively; no additional DRAQ7-positive cells were observed during the extended period. Changes in cell position and movement within the hydrogel were observed in all eight longitudinally evaluated extraction experiments. These findings indicate that although direct micropipette contact may cause localized damage to the aspirated cell, the extraction procedure can preserve membrane integrity in the remaining cells of the recovered multicellular cluster, and that cell division may occur after extraction.

**Figure 4.**
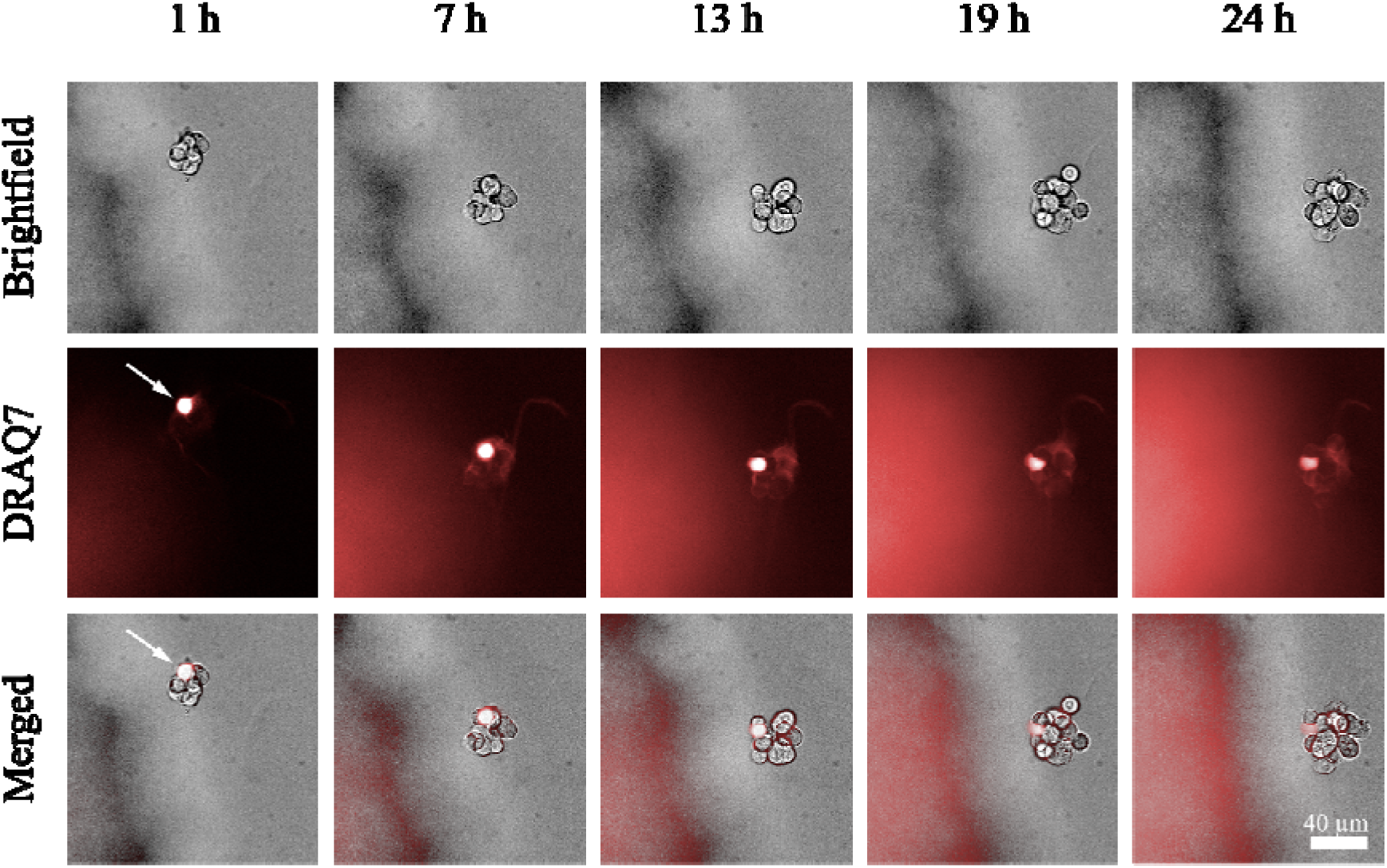
Longitudinal monitoring of an extracted multicellular cluster following micropipette-based manipulation. (extraction *20260803_H_M_02_02*). After removing the cells, DRAQ7 staining was added to the cell culture medium to label the nuclei of membrane-compromised cells. Representative brightfield images (top row), Alexa 647 fluorescence images (middle row), and merged brightfield/fluorescence images (bottom row) were acquired at 1, 7, 13, 19, and 24-h after the start of imaging, showing the post-extraction morphology and fluorescence signal of the recovered cell cluster. The white arrow indicates the cell that was directly aspirated by the micropipette during extraction and was consequently damaged, resulting in a persistent DRAQ7-positive signal. The scale bar represents 40 µm.

In both single-cell extractions evaluated with DRAQ7 (*20260917_T_S_01_01* and *20260917_T_S_01_02*), the isolated cell was DRAQ7-positive from the first imaging time point. In the doublet extraction *20260901_T_D_01_01*, one of the two recovered cells became DRAQ7-positive after approximately 5-h, whereas the other remained DRAQ7-negative throughout the observation window. These observations suggest that, under the present operating conditions, extraction of viable multicellular clusters is more robust than isolation of individual viable cells. The corresponding single-cell extraction experiments and outcomes are documented in **Supplementary Note 1** and the **Supplementary Videos**.

Post-extraction time-lapse annotations were evaluated for eight extraction experiments (four HeLa Kyoto and four T-47D) to analyse the morphological changes of the extracted cells. Twelve individual cell contours were followed in seven experiments; in the remaining HeLa Kyoto experiment *20260803_H_M_02_02*, individual boundaries could not be reliably distinguished, so the extracted unit was annotated as a whole cluster. In experiment *20260925_H_M_01_01*, the extracted cells separated during manipulation, one DRAQ7-positive cell was completely disrupted and therefore excluded from annotation, while the remaining DRAQ7-negative cell was annotated. The two single-cell extractions from specimen *20260917_T_01* were followed in separate Operetta fields. Between the first and final frames, the annotated focal-plane area decreased in five individual cells and increased in seven. The whole-cluster contour increased from 1058.4 to 2568.9 µm². With the specimen immobilized in hydrogel, net XY centroid displacement ranged from 7.6 to 87.6 µm among individual cells and was 99.3 µm for the whole-cluster contour. The corresponding signed changes in absolute focus-plane position are reported in Table 1. Individual-cell contours showed heterogeneous changes in area and circularity, whereas the whole-cluster contour expanded and became less circular. Abrupt focus-plane shifts were confined to the extracted cells, whereas the parent spheroid and surrounding non-extracted objects remained in a stable focal plane.

**Table 1.** Focal-plane morphology and displacement during post-extraction observation. Initial and final measurements correspond to the first and last acquisition frames. The fields from specimen *20260917_T_01* represent two separate, sequential single-cell extraction experiments. Area and circularity are medians across annotated cells at the first and final frames; XY displacement is given as a range and Z focus change as a median (see Methods). Z focus change is not a reconstructed 3D displacement. ^a^ Last frame at 23-h instead of 24-h. * Cell division was observed in this extracted cluster during the observation period.

| Extraction ID | Cell line | Extracted unit | Annotated objects | Area ($\mu\text{m}^2$) | Circularity | XY displacement ($\mu\text{m}$) | Z focus change ($\mu\text{m}$) | DRAQ7 □ cells at last frame (n) |
| --- | --- | --- | --- | --- | --- | --- | --- | --- |
| *20260803_H_M_02_02* | HeLa Kyoto | cluster | whole cluster | 1058.4 → 2568.9* | 0.719 → 0.426 | 99.3 | -10.6 | 1 |
| *20260812_H_M_01_01* | HeLa Kyoto | cluster | 3 cells | 72.9 → 199.6 | 0.948 → 0.952 | 38.5-42.6 | -199.0 | 1 |
| *20260817_H_M_01_04* | HeLa Kyoto | cluster | 2 cells | 148.2 → 161.4 | 0.942 → 0.952 | 21.1-24.8 | -77.8 | 1 |
| *20260925_H_M_01_01* | HeLa Kyoto | cluster | 1 cell | 210.2 → 241.9 | 0.959 → 0.951 | 37.8 | -159.5 | 0 |
| *20260901_T_D_01_01* | T-47D | doublet | 2 cells | 223.8 → 171.1 | 0.927 → 0.970 | 25.1-87.6 | -238.2 | 1 |
| *20260904_T_D_01_02* | T-47D | doublet | 2 cells | 194.5 → 215.3 | 0.967 → 0.955 | 7.6-12.1 | +4.2 | 0 |
| 20260917_T<br>_S_01_01<sup>a</sup> | T-47D | single cell | 1 cell | 329.9 →<br>283.1 | 0.701 →<br>0.786 | 22.1 | -93.4 | 1 |
| 20260917_T<br>_S_01_02<sup>a</sup> | T-47D | single cell | 1 cell | 262.3 →<br>373.2 | 0.751 →<br>0.797 | 11.0 | -131.7 | 1 |

## 4. Discussion

3D cell cultures have gained attention in the last couple of years, indicating that such models may take important roles in drug discovery, development studies, and that we are heading to the organ-on-demand era. Spheroids, as one of the most basic but extensively used 3D models, highlight that there are still areas that we have to understand better to move forward to more complex models. Over the years, the community has realised that the inconsistency in reporting experimental parameters and the high variability in generation methods can cause discrepancies^32^. Furthermore, we tend to accept that generating the same spheroids with similar parameters is not that simple^4,33^. Nonetheless, to see the inner parts of the relatively large samples requires time-consuming fixation and optical clearing techniques that prevent further live cell investigation^34,35^. These works highlighted that even though many publications report work with spheroids, our options to investigate the spheroids are limited.

To gain in-depth information, there are methods to destroy the 3D structure of spheroids either physically or enzymatically, however, these methods only allow a single time point without the option to select the target cells^36^. However, most recent publications highlight the need for real-time analysis of 3D cell cultures^37^.

Cell migration has become a major research focus, since in a 3D environment cells must navigate dense and heterogeneous extracellular matrices, squeeze through narrow spaces, and dynamically adapt their migration. Since the 3D migration of cells leaves many open questions in tissue development, immune responses, tissue repair, and cancer invasion, researchers increasingly recognize the need for high-precision tools enabling precise manipulation and monitoring^38,39^. Beyond migration, cell morphology and molecular dynamics play a crucial role in uncovering disease-related morpho-molecular relationships. Together with the spatial information, they reveal how cells reshape themselves to fulfil specific functions and help identify the mechanisms driving abnormal cellular behaviors in pathological conditions^40,41^. Although methods exist to acquire molecular and spatial information from intact 3D cell cultures, they are limited by fixation, sample size and time-consuming procedures^42,43^. Therefore, the ability to locally extract and reposition viable multicellular clusters from intact 3D cultures may support several experimental applications where conventional bulk dissociation is difficult to implement.

In this work, we propose an alternative method to study dense tumor spheroids by using a pressure-controlled micropipette-based extraction. Through a series of optimisations, the safe extraction of cell clusters from the outer layers of spheroids embedded in hydrogel became accessible in a reliable fashion. First, we systematically tested the hydrogel component to embed and secure the position of the spheroid while allowing the navigation of the glass capillary. Then, we tested several pressure settings that were adjusted to the viscosity of the hydrogel to move freely without the hydrogel adhering to the inner wall of the glass capillary, causing blockage and preventing proper pressure transmission. Finally, we tested a range of pipette tip sizes and made conclusions about the tip diameter compared to the cell itself to be able to maintain the right negative pressure or attaching force. Combining such knowledge, targeted cells can be removed from the outer layers and repositioned freely within the hydrogel. Transfer to a separate holder is not performed directly, but the deposited cluster can be excised together with the surrounding hydrogel and released by enzymatic degradation of the matrix, allowing the isolated cells to be recovered for downstream analysis.

These removed cell clusters were monitored for 24-h and showed evidence of retained viability after the extraction. No DRAQ7-positive signal was observed near the apparent extraction region of the parent spheroid immediately after DRAQ7 screening, although the exact extraction location, particularly along the Z-axis, could not be reliably determined. Besides, subsequent cell division was observed in one of the six multicellular-cluster extraction experiments evaluated, providing functional evidence that some of the recovered multicellular material retained proliferative capacity. The limited frequency of observed division may partly reflect the small size of the recovered clusters (approximately 2-15 cells), which may provide reduced cell–cell signalling and paracrine support required for sustained proliferative activity. However, this remains speculative because the effect of cluster size on proliferation was not directly investigated. Cell movement was also observed during longitudinal imaging, although movement alone was not considered an independent criterion of viability. The preservation of membrane integrity and the observation of cell division in one experiment support the feasibility of continued post-extraction cultivation and longitudinal observation. The platform is intended for future applications at both the multicellular-cluster and single-cell levels, including controlled cell migration studies, donor–acceptor experiments, drug-response analysis, studies using organoid models, and downstream molecular profiling.

One application is the longitudinal investigation of cell migration after spatially controlled removal from a spheroid. Extracted clusters can be positioned at defined distances from the parent spheroid and monitored over time to determine whether individual cells remain within the recovered cluster, migrate into the surrounding matrix, or move back towards the original source. Such experiments could be used to quantify migration directionality, velocity, persistence, and the influence of local matrix conditions on collective and individual cell behaviour. Localized recovery of cellular material may also facilitate downstream molecular analysis. In contrast to complete spheroid dissociation, the proposed method preserves information about the original sampling position and leaves the remaining spheroid available for further cultivation, imaging, or repeated intervention. The extracted clusters could be analysed using omic methods. Combining repeated localized extraction with time-resolved imaging could establish a link between molecular profiles and dynamic cellular behaviours such as proliferation, migration, or treatment response.

The method presented in this study serves as an early proof-of-concept, demonstrating that the removal of viable cell clusters is feasible in a reliable fashion. While this initial showcase highlights the potential of the approach, there remains substantial room for refinement and optimization in future developments. The cell extraction was demonstrated on spheroids generated from well-known cell lines; however, the same principle may be applicable to many other cell lines or more complex 3D models, including patient-derived organoids and engineered organotypic cultures. On the other hand, 3D samples with strong cell-cell adhesion, high cellular density, or advanced maturation will require additional model-specific testing and careful parameter optimization to ensure reliable performance. Although the common aim is to extract living single-cells from a dense spheroid, the reliable extraction will require refinement of micropipette tip geometry, pressure control, local enzymatic treatment, and release conditions. Nevertheless, while attaching to and isolating an individual cell from the spheroid was feasible, the integrity of the cell membrane was likely compromised during either the extraction or the release step. In our experiments this damage was restricted to the cell in direct contact with the micropipette, suggesting a localised mechanical origin rather than an intrinsic limitation of the approach. Optimisation of the aspiration pressure profile and of the tip geometry relative to cell size is therefore expected to mitigate this effect, and is a subject of our ongoing development.

Instead of focusing on a single-cell, we suggest removing small cell clusters (2-15 cells), where the cell directly contacted or aspirated by the micropipette is particularly susceptible to membrane damage, whereas neighbouring cells within the same extracted cluster can remain viable and functionally uncompromised. Accessing deeper layers of the sample introduces further challenges. Navigating through the surrounding matrix and the spheroid itself increases mechanical resistance and the risk of damaging both the target cell and the parent culture. However, working inside a dense 3D structure requires precision where high-resolution visibility is essential. The current depth range, however, is limited primarily by imaging rather than by the manipulator itself: with the widefield optics used here, neither the capillary tip nor the target cell can be reliably resolved below the outermost cell layers. Extending the approach to deeper layers will therefore require combining depth-resolved imaging with a sample geometry that remains physically accessible to the capillary, and with a means of visualising the capillary tip within the same imaging modality. The current workflow is manual and depends on operator judgement for target selection, micropipette positioning, pressure adjustment, extraction, and deposition of the recovered material. This limits throughput and may introduce operator-dependent variability. Future automation could combine image-based detection of spheroid boundaries and target regions, computer-controlled micromanipulator trajectories, adaptive pressure regulation, and automated tracking of extracted cells or clusters.

Overall, the present work establishes an initial methodology for localized extraction and manipulation of viable multicellular clusters from the surface of living 3D cultures. The present proof-of-concept demonstrates that multicellular clusters can be detached from the outer layers of hydrogel-embedded spheroids, displaced within the surrounding matrix, and subsequently relocated on a high-content imaging system. Such controlled positioning could support single-cell printing, generation of defined mixed-cell arrangements, and investigation of how the initial position of an individual cell affects subsequent proliferation, migration, differentiation, or integration into a multicellular system. Compared with complete enzymatic or mechanical dissociation, the presented approach provides localized access to selected regions while preserving the parent spheroid as a continuous 3D structure. The method is not intended to replace bulk dissociation when high cellular yield is the primary objective. Instead, it represents a complementary strategy for experiments in which spatial origin, continued cultivation, repeated sampling, and controlled manipulation are more important than throughput. Future work will focus on reducing direct-contact damage and achieving reliable extraction of viable single cells from deeper regions of 3D cultures, and introducing semi-automated and automated control, opening the door to high-fidelity molecular, morphological, and spatial profiling for complex biological models.

## Supporting information

Supplementary_data_1_Logsheets

Supplementary_data_2_metrics

Supplementary_Softwares

## Data availability

The attempt-level experiment outcome log is provided in **Supplementary Note 1**. The cell-level measurements underlying Table 1 are provided in **Supplementary Data 2**, and detailed experimental log sheets for the eight DRAQ7-evaluated extractions are provided in **Supplementary Data 1**. The Operetta post-extraction screening datasets, corresponding image annotations, and multichannel TIFF files are available in **Figshare** at https://doi.org/10.6084/m9.figshare.c.8785665. Representative extraction and longitudinal-imaging records are provided as **Supplementary Videos**.

## Declaration of generative AI and AI-assisted technologies in the manuscript preparation process

During the preparation of this work the authors used Astra (OpenAI) and Opus 5.5 (Anthropic) to increase readability and grammar check of the text. After using these tool services, the authors reviewed and edited the content as needed and took full responsibility for the content of the published article.

## Code availability

The custom *Python* script used to calculate and summarize the image-annotation measurements is provided as **Supplementary Software 1**. The custom script used to generate Figure 2 from the extraction log is provided as **Supplementary Software 2**.

## Acknowledgements

The authors acknowledge support from the TKP2021-EGA09 (TKP-31-8/PALY-2021), TKCS-2024/73, Horizon-BIALYMPH (2023-1.2.1-ERA_NET-2023-00002), Horizon - SYMMETRY (2019-2.1.7-ERA-NET-2022-00044), Horizon-RIA - SWEEPICS (101135053), Horizon 2020 - FAIR CHARM (101016457), HAS-NAP3 (NAP2022-I-6/2022), PerMel-AI (EPPERMED2024−590), OTKA-SNN_21 (139455/ARRS) and NKKP (ADVANCED_24_151202), 2026-4.1.1-MISSZIÓK-2026-00025. M.S. and F.P. acknowledge support from the MAECI Science and Technology Cooperation Italy-South Korea Grant Years 2026–2028 by the Italian Ministry of Foreign Affairs and International Cooperation (CUP project: J53C26000240006). M.H. acknowledges support from the János Bolyai Research Scholarship of the Hungarian Academy of Sciences and the NKKP (STARTING_25_152376).

## Author contributions statement

Conceptualization: V.P., M.M., M.S., F.P., A.D., P.H.

Data curation: V.P., M.M., M.S., N.H.

Formal analysis: V.P., M.M., M.S., N.H., M.H., A.D.

Funding acquisition: F.P., P.H.

Investigation: V.P., M.M., M.S., N.H., M.H., A.D.

Methodology: V.P., M.M., M.S., N.H., M.H., A.D.

Project administration: F.P., P.H.

Resources: P.H.

Software: V.P., M.M.

Supervision: M.H., F.P., A.D., P.H.

Validation: V.P., M.M., M.S., N.H.

Visualization: V.P., M.M.

Writing – original draft: V.P., M.M., M.S., N.H., F.P.

Writing – review & editing: M.H., A.D., P.H.

