## Supplementary_data_1_Logsheets for "Spheroid surgery: a first-generation micromanipulation platform for extracting viable cell clusters from the outer layers of hydrogel-embedded spheroids": 20260803_H_M_02_02.pdf

### Single-Cell Extraction — Experiment Log

20260803_H_M_02

#### 01 Identification & metadata

| | |
|---|---|
| Date | 2026-08-03 |
| Operator | Viktor |
| Sample ID | 02 |
| Environment | H — Hydrogel |
| Configuration | M — Multiple / cluster |

#### 02 Cell line & spheroid

| | |
|---|---|
| Cell line | HeLa Kyoto |
| Passage number | 3/2/5 |
| Spheroid age | 3 days |
| Spheroid preparation | — |

#### 03 Hydrogel — TrueGel3D

##### COMPONENTS PIPETTED — PER GEL

| COMPONENT | VOLUME (ML) | NOTE |
|---|---|---|
| SLO-Dextran stock | 0.9 | |
| CD crosslinker | 1.35 | First open |
| Water | 8.55 | |
| Buffer | 1.2 | |
| Cell suspension | 3 | |

| | |
|---|---|
| Target final volume | 15 µL |

##### CROSSLINKER ALIQUOT & THIOL OXIDATION

| | |
|---|---|
| Times this aliquot was opened | 1 |
| Other deviations from protocol | First open of the crosslinker |

#### 04 Dissociation / enzyme

| | |
|---|---|
| Enzyme used | Trypsin |

#### 05 Micropipette & pressures

##### TIP GEOMETRY

| | |
|---|---|
| Tip diameter | 4.5 µm |

##### PRESSURE STEPS

| STEP | PRESSURE (MBAR) | POLARITY | DURATION (S) | NOTE |
|---|---|---|---|---|
| Approach | 300 | positive | | keeps tip clear while entering gel |
| Extraction | -500 | negative | | aspiration of the target cell |
| Deposition | 300 | positive | | expel at target site — remove row if unused |

| | |
|---|---|
| Pressure controller | Custom |
| Compensation pressure held? | No |
| Compensation pressure | — |

#### 06 Extraction run

| | |
|---|---|
| Extraction time | 3.5 min |
| Attempts | 2 |
| Cells recovered | 14 |
| Visible tip clogging? | No |
| Rig / instrument | Luigs & Neumann LNscope (upright) + LN Junior Compact micromanipulator |
| Run observations | — |

#### 07 Imaging plate

| | |
|---|---|
| Plate type | ibidi µ-Slide 1 Well Glass Bottom (82107) |
| Substrate | 170 µm borosilicate, RI 1.523 |

#### 09 Imaging

| | |
|---|---|
| Viability stain | DRAQ7 (Thermo D15105) |
| DRAQ7 concentration | 3 µM |

#### 10 Outcome

| | |
|---|---|
| Successful extraction? | Yes |
| Membrane intact at deployment? | Yes |
| Division observed? | Yes |
| Time to first division | 6 h |
| Time to DRAQ7-positive | — |
| Final cell count | — |
| Outcome notes | — |

#### 11 Notes / observations

We extracted a small chunk of cells. When we extracted we lost the attachment for a short period, but after the re-approach we were able to move the cells to a relatively big distance from the original spheroid (higher around 200 µm), The only cell what died was the cell we attached the pipette. Some cells started to divide after 6 h. After the 24h screening we put the sample to the incubator and checked daily.
