## Supplementary_data_1_Logsheets for "Spheroid surgery: a first-generation micromanipulation platform for extracting viable cell clusters from the outer layers of hydrogel-embedded spheroids": 20260812_H_M_01_01.pdf

### Single-Cell Extraction — Experiment Log

20260812_H_M_01

#### 01 Identification & metadata

| | |
|---|---|
| Date | 2026-08-12 |
| Operator | Viktor |
| Sample ID | 01 |
| Environment | H — Hydrogel |
| Configuration | M — Multiple / cluster |

#### 02 Cell line & spheroid

| | |
|---|---|
| Cell line | HeLa Kyoto |
| Passage number | 3/2/8 |
| Spheroid age | 2 days |
| Spheroid preparation | — |

#### 03 Hydrogel — TrueGel3D

COMPONENTS PIPETTED — PER GEL

| COMPONENT | VOLUME (ML) | NOTE |
|---|---|---|
| SLO-Dextran stock | 0.9 | |
| CD crosslinker | 1.35 | 3. open |
| Water | 8.55 | |
| Buffer | 1.2 | |
| Cell suspension | 3 | |

| | |
|---|---|
| Target final volume | 15 µL |

CROSSLINKER ALIQUOT & THIOL OXIDATION

| | |
|---|---|
| Times this aliquot was opened | 3 |
| Other deviations from protocol | In the hydrogel we had a lot of bubbles under the hydrogel |

#### 04 Dissociation / enzyme

| | |
|---|---|
| Enzyme used | Trypsin |

#### 05 Micropipette & pressures

TIP GEOMETRY

| | |
|---|---|
| Tip diameter | 4.5 µm |

PRESSURE STEPS

| STEP | PRESSURE (MBAR) | POLARITY | DURATION (S) | NOTE |
|---|---|---|---|---|
| Approach | 200 | positive | | keeps tip clear while entering gel |
| Extraction | -500 | negative | | aspiration of the target cell |
| Deposition | 200 | positive | | expel at target site — remove row if unused |

| | |
|---|---|
| Pressure controller | Custom |
| Compensation pressure held? | No |
| Compensation pressure | — |

#### 06 Extraction run

| | |
|---|---|
| Extraction time | — |
| Attempts | 2 re-approach |
| Cells recovered | 3 |
| Visible tip clogging? | No |
| Rig / instrument | Luigs & Neumann LNscope (upright) + LN Junior Compact micromanipulator |
| Run observations | We extracted approximately 4 cells, 3 is visible under the Operetta screen |

#### 07 Imaging plate

| | |
|---|---|
| Plate type | ibidi µ-Slide 1 Well Glass Bottom (82107) |
| Substrate | 170 µm borosilicate, RI 1.523 |

#### 09 Imaging

| | |
|---|---|
| Viability stain | DRAQ7 (Thermo D15105) |
| DRAQ7 concentration | 3 µM |

#### 10 Outcome

| | |
|---|---|
| Successful extraction? | Yes |
| Membrane intact at deployment? | Yes |
| Division observed? | No |
| Time to first division | — |
| Time to DRAQ7-positive | — |
| Final cell count | 3 |
| Outcome notes | — |

#### 11 Notes / observations

One cells give DRAQ7 signal but the other two not
