## Supplementary_data_1_Logsheets for "Spheroid surgery: a first-generation micromanipulation platform for extracting viable cell clusters from the outer layers of hydrogel-embedded spheroids": 20260817_H_M_01_04.pdf

### Single-Cell Extraction — Experiment Log

20260817_H_M_01

#### 01 Identification & metadata

| | |
|---|---|
| Date | 2026-08-17 |
| Operator | Viktor |
| Sample ID | 01 |
| Environment | H — Hydrogel |
| Configuration | M — Multiple / cluster |

#### 02 Cell line & spheroid

| | |
|---|---|
| Cell line | HeLa Kyoto |
| Passage number | 3/2/9 |
| Spheroid age | 3 days |
| Spheroid preparation | — |

#### 03 Hydrogel — TrueGel3D

##### COMPONENTS PIPETTED — PER GEL

| COMPONENT | VOLUME (ML) | NOTE |
|---|---|---|
| SLO-Dextran stock | 0.9 | |
| CD crosslinker | 1.35 | 5. open |
| Water | 8.55 | |
| Buffer | 1.2 | |
| Cell suspension | 3 | |

| | |
|---|---|
| Target final volume | 15 µL |

##### CROSSLINKER ALIQUOT & THIOL OXIDATION

| | |
|---|---|
| Times this aliquot was opened | 5 |
| Other deviations from protocol | The hydrogel density was perfect, after the 3. hour the cells sinked a littlebit. |

#### 04 Dissociation / enzyme

| | |
|---|---|
| Enzyme used | Trypsin |

#### 05 Micropipette & pressures

##### TIP GEOMETRY

| | |
|---|---|
| Tip diameter | 4 µm |

##### PRESSURE STEPS

| STEP | PRESSURE (MBAR) | POLARITY | DURATION (S) | NOTE |
|---|---|---|---|---|
| Approach | 200 | positive | | keeps tip clear while entering gel |
| Extraction | -500 | negative | | aspiration of the target cell |
| Deposition | 200 | positive | | expel at target site — remove row if unused |

| | |
|---|---|
| Pressure controller | Custom |
| Compensation pressure held? | No |
| Compensation pressure | — |

#### 06 Extraction run

| | |
|---|---|
| Extraction time | 5 min |
| Attempts | 4 |
| Cells recovered | 1 |
| Visible tip clogging? | No |
| Rig / instrument | Luigs & Neumann LNscope (upright) + LN Junior Compact micromanipulator |
| Run observations | We extracted 3 cells but two of them died right after or during the extraction. The 4. cell just deattached during the extraction. |

#### 07 Imaging plate

| | |
|---|---|
| Plate type | ibidi µ-Slide 1 Well Glass Bottom (82107) |
| Substrate | 170 µm borosilicate, RI 1.523 |

#### 09 Imaging

| | |
|---|---|
| Viability stain | DRAQ7 (Thermo D15105) |
| DRAQ7 concentration | 3 µM |

#### 10 Outcome

| | |
|---|---|
| Successful extraction? | Yes |
| Membrane intact at deployment? | Yes |
| Division observed? | No |
| Time to first division | — |
| Time to DRAQ7-positive | — |
| Final cell count | 2 |
| Outcome notes | — |

#### 11 Notes / observations

From the 4 extracted cells only one remained DRAQ7 negative after the extraction but it remained DRAQ7 negative during the 24 hour screening, and we tried an additional 24 hour and it was still DRAQ7 negative in the end.
