## Supplementary_data_2_metrics for "Spheroid surgery: a first-generation micromanipulation platform for extracting viable cell clusters from the outer layers of hydrogel-embedded spheroids": Table1.docx

| **Extraction ID** | **Cell line** | **Target** | **Annotated objects** | **Area (µm²)** | **Circularity** | **XY displacement (µm)** | **Z focus change (µm)** | **DRAQ7⁺ cells at last frame (n)** | **Division** |
| --- | --- | --- | --- | --- | --- | --- | --- | --- | --- |
| 20260803_H_M_02_02 | HeLa Kyoto | cluster | whole cluster | 1058.4 → 2568.9 | 0.719 → 0.426 | 99.3 | -10.6 | 1 | + |
| 20260812_H_M_01_01 | HeLa Kyoto | cluster | 3 cells | 72.9 → 199.6 | 0.948 → 0.952 | 38.5–42.6 | -199.0 | 1 | - |
| 20260817_H_M_01_04 | HeLa Kyoto | cluster | 2 cells | 148.2 → 161.4 | 0.942 → 0.952 | 21.1–24.8 | -77.8 | 1 | - |
| 20260925_H_M_01_01 | HeLa Kyoto | cluster | 1 cell | 210.2 → 241.9 | 0.959 → 0.951 | 37.8 | -159.5 | 0 | - |
| 20260901_T_D_01_01 | T-47D | doublet | 2 cells | 223.8 → 171.1 | 0.927 → 0.970 | 25.1–87.6 | -238.2 | 1 | - |
| 20260904_T_D_01_02 | T-47D | doublet | 2 cells | 194.5 → 215.3 | 0.967 → 0.955 | 7.6–12.1 | +4.2 | 0 | - |
| 20260917_T_S_01_01ᵃ | T-47D | single cell | 1 cell | 329.9 → 283.1 | 0.701 → 0.786 | 22.1 | -93.4 | 1 | - |
| 20260917_T_S_01_02ᵃ | T-47D | single cell | 1 cell | 262.3 → 373.2 | 0.751 → 0.797 | 11.0 | -131.7 | 1 | - |

ᵃ Last frame at 23 h instead of 24 h.
