## Supplementary_Softwares for "Spheroid surgery: a first-generation micromanipulation platform for extracting viable cell clusters from the outer layers of hydrogel-embedded spheroids": make_table1.py

```
"""Build Table 1 from per-object metrics (output of measure_annotations.py).

Usage:
    python make_table1.py [--units table1_units.csv] [--metrics metrics_dir] [--out out_dir]

The units CSV maps each extraction ID to its metrics file prefix and holds the
per-extraction observations taken from the experiment log (DRAQ7-positive
cells at the last frame, division).
"""
import argparse
from pathlib import Path

import numpy as np
import pandas as pd
from docx import Document
from docx.enum.section import WD_ORIENT
from docx.shared import Pt

DATA = Path(__file__).resolve().parent.parent / 'Supplementary_Data_1'
REQUIRED = ['unit_id', 'cell_line', 'target', 'metrics_prefix', 'draq7_positive_final', 'division']
ARROW = ' → '


def parse_args():
    p = argparse.ArgumentParser(description=__doc__)
    p.add_argument('--units', type=Path, default=DATA / 'table1_units.csv')
    p.add_argument('--metrics', type=Path, default=DATA / 'metrics')
    p.add_argument('--out', type=Path, default=DATA / 'table1')
    p.add_argument('--pattern', default='**/*_metrics.csv')
    p.add_argument('--endpoints', default='23,24', help='Accepted last timepoints, h')
    p.add_argument('--digits', type=int, default=1, help='Decimals for µm and µm² values')
    p.add_argument('--circ-digits', type=int, default=3)
    return p.parse_args()


def load_units(path):
    u = pd.read_csv(path, dtype=str).fillna('')
    missing = [c for c in REQUIRED if c not in u]
    if missing:
        raise ValueError(f'{path.name}: missing columns {missing}')
    if u.unit_id.duplicated().any():
        raise ValueError(f'Duplicate unit_id: {u.unit_id[u.unit_id.duplicated()].tolist()}')
    u['draq7_positive_final'] = u.draq7_positive_final.astype(int)
    return u


def load_metrics(prefix, args):
    hits = [p for p in args.metrics.glob(args.pattern) if p.name.startswith(prefix)]
    if len(hits) != 1:
        raise ValueError(f'Expected one metrics file for {prefix}, found {hits}')
    d = pd.read_csv(hits[0])
    if 'focus_absolute_um' not in d:
        raise ValueError(f'{hits[0].name}: no focus_absolute_um; re-run measure_annotations.py')
    return d


def summarize(unit, d, endpoints):
    end_t = int(d.timepoint.max())
    if d.timepoint.min() != 0 or end_t not in endpoints:
        raise ValueError(f'{unit.unit_id}: unexpected time range {d.timepoint.min()}-{end_t}')
    objs = []
    for label, tr in d.groupby('label'):
        if sorted(tr.timepoint) != list(range(end_t + 1)):
            raise ValueError(f'{unit.unit_id}: incomplete track for label {label}')
        np.testing.assert_allclose(tr.circularity, 4 * np.pi * tr.area_um2 / tr.perimeter_um ** 2,
                                   rtol=1e-10, atol=1e-9)
        a = tr[tr.timepoint == 0].iloc[0]
        b = tr[tr.timepoint == end_t].iloc[0]
        objs.append(dict(unit_id=unit.unit_id, source=a.source, label=int(label), endpoint_h=end_t,
                         initial_area_um2=a.area_um2, final_area_um2=b.area_um2,
                         initial_circularity=a.circularity, final_circularity=b.circularity,
                         xy_displacement_um=float(np.hypot(b.centroid_x_um - a.centroid_x_um,
                                                           b.centroid_y_um - a.centroid_y_um)),
                         z_focus_displacement_um=float(b.focus_absolute_um - a.focus_absolute_um)))
    o = pd.DataFrame(objs)
    first, last = d[d.timepoint == 0], d[d.timepoint == end_t]
    row = dict(unit_id=unit.unit_id, cell_line=unit.cell_line, target=unit.target,
               source=d.source.iloc[0], endpoint_h=end_t,
               annotation_level=first.annotation_level.iloc[0], n_annotated_objects=len(o),
               initial_median_area_um2=float(first.area_um2.median()),
               final_median_area_um2=float(last.area_um2.median()),
               initial_median_circularity=float(first.circularity.median()),
               final_median_circularity=float(last.circularity.median()),
               min_xy_displacement_um=o.xy_displacement_um.min(),
               max_xy_displacement_um=o.xy_displacement_um.max(),
               median_z_focus_displacement_um=float(o.z_focus_displacement_um.median()),
               draq7_positive_cells_at_final_frame=unit.draq7_positive_final,
               division_observed=unit.division)
    return row, objs


def display_table(n, args, max_end):
    f, c = f'.{args.digits}f', f'.{args.circ_digits}f'

    def objects(r):
        if r.annotation_level == 'cluster':
            return 'whole cluster'
        return f'{r.n_annotated_objects} cell' + ('s' if r.n_annotated_objects > 1 else '')

    def xy(r):
        lo, hi = format(r.min_xy_displacement_um, f), format(r.max_xy_displacement_um, f)
        return lo if r.n_annotated_objects == 1 else f'{lo}–{hi}'

    def uid(r):
        return r.unit_id + ('ᵃ' if r.endpoint_h != max_end else '')

    return pd.DataFrame({
        'Extraction ID': [uid(r) for r in n.itertuples()],
        'Cell line': n.cell_line,
        'Target': n.target,
        'Annotated objects': [objects(r) for r in n.itertuples()],
        'Area (µm²)': [format(r.initial_median_area_um2, f) + ARROW + format(r.final_median_area_um2, f)
                       for r in n.itertuples()],
        'Circularity': [format(r.initial_median_circularity, c) + ARROW + format(r.final_median_circularity, c)
                        for r in n.itertuples()],
        'XY displacement (µm)': [xy(r) for r in n.itertuples()],
        'Z focus change (µm)': [format(r.median_z_focus_displacement_um, '+' + f) for r in n.itertuples()],
        'DRAQ7⁺ cells at last frame (n)': n.draq7_positive_cells_at_final_frame.astype(str),
        'Division': n.division_observed,
    })


def write_docx(table, path, max_end, short_ends):
    doc = Document()
    sec = doc.sections[0]
    sec.orientation = WD_ORIENT.LANDSCAPE
    sec.page_width, sec.page_height = sec.page_height, sec.page_width
    t = doc.add_table(rows=1, cols=len(table.columns))
    t.style = 'Table Grid'
    for cell, name in zip(t.rows[0].cells, table.columns):
        cell.text = name
        cell.paragraphs[0].runs[0].bold = True
    for row in table.itertuples(index=False):
        for cell, v in zip(t.add_row().cells, row):
            cell.text = str(v)
    for r in t.rows:
        for cell in r.cells:
            for par in cell.paragraphs:
                for run in par.runs:
                    run.font.size = Pt(8)
    if short_ends:
        doc.add_paragraph(f'ᵃ Last frame at {", ".join(map(str, short_ends))} h instead of {max_end} h.')
    doc.save(path)


def main():
    args = parse_args()
    endpoints = {int(x) for x in args.endpoints.split(',')}
    units = load_units(args.units)
    rows, objs = [], []
    for u in units.itertuples():
        r, o = summarize(u, load_metrics(u.metrics_prefix, args), endpoints)
        rows.append(r)
        objs.extend(o)
    n = pd.DataFrame(rows)
    max_end = int(n.endpoint_h.max())
    short_ends = sorted(set(n.endpoint_h[n.endpoint_h != max_end]))
    table = display_table(n, args, max_end)

    args.out.mkdir(parents=True, exist_ok=True)
    n.to_csv(args.out / 'Table1_numeric.csv', index=False)
    pd.DataFrame(objs).to_csv(args.out / 'Table1_individual_objects.csv', index=False)
    table.to_csv(args.out / 'Table1.csv', index=False, encoding='utf-8-sig')
    write_docx(table, args.out / 'Table1.docx', max_end, short_ends)
    print(table.to_string(index=False))
    specimens = n.unit_id.str.rsplit('_', n=1).str[0].str.replace(r'_[A-Z]_(\d+)$', r'_\1', regex=True).nunique()
    print(f'\n{len(n)} extractions, {specimens} specimens, '
          f'{int((n.annotation_level == "cell").mul(n.n_annotated_objects).sum())} annotated cells')


if __name__ == '__main__':
    main()
```
