## Supplementary_Softwares for "Spheroid surgery: a first-generation micromanipulation platform for extracting viable cell clusters from the outer layers of hydrogel-embedded spheroids": measure_annotations.py

```
"""Per-object morphology and focal-plane position from napari label masks.

Usage:
    python measure_annotations.py config.json

Each entry of "annotation_dirs" in the config is one Operetta measurement
folder holding label masks (one in-focus plane annotated per object and
timepoint). Pixel size, Z positions and acquisition times are read from the
measurement's Operetta Index.xml, found by walking up from the mask folder.
One <mask>_metrics.csv is written per mask file, plus ALL_metrics.csv.

Z values describe the selected focal plane (absolute stage position from
the metadata), not a reconstructed 3D trajectory.
"""
import argparse
import json
import re
import warnings
import xml.etree.ElementTree as ET
from collections import defaultdict
from pathlib import Path

import numpy as np
import pandas as pd
import tifffile
from skimage.measure import regionprops_table

DEFAULTS = {
    "annotation_dirs": [],
    "mask_pattern": "*.tif",
    "output_dir": "metrics",
    # Meaning of the leading axis of a 3-axis mask: "Z" (one timepoint,
    # objects on their own focus planes) or "T" (time series of 2D masks).
    "ambiguous_axis": "Z",
    "mask_axes_override": None,          # e.g. "TZYX"
    # Regex with one group capturing the timepoint when each file is one
    # timepoint of a series, e.g. "_t(\\d+)"; null = one file per series.
    "timepoint_pattern": None,
    "series_name": None,
    "generic_folder_names": ["annotations", "assembled", "images", "metrics"],
    "index_xml_path": None,
    "index_xml_names": ["index.xml", "index.idx.xml"],
    "index_search_depth": 4,
    "brightfield_channel_keyword": "brightfield",
    "time_interval_h": 1.0,
    # Regexes matched against mask file names; matching files are whole-
    # cluster outlines and are never pooled with single cells.
    "cluster_level_patterns": [],
    "z_jump_threshold_um": 8.0,          # QC only, not a biological threshold
    "min_instance_area_px": 0,
    "background_label": 0,
    "overwrite": True,
}

BASE_PROPERTIES = (
    "label", "area", "perimeter", "centroid", "axis_major_length",
    "axis_minor_length", "eccentricity", "solidity", "extent",
    "orientation", "equivalent_diameter_area",
)

OUTPUT_COLUMNS = [
    "source", "annotation_level", "n_labels_in_source", "timepoint", "time_h", "label",
    "area_um2", "perimeter_um", "equivalent_diameter_um", "axis_major_um", "axis_minor_um",
    "aspect_ratio", "circularity", "eccentricity", "solidity", "extent", "orientation_deg",
    "centroid_x_um", "centroid_y_um", "step_um", "displacement_from_origin_um",
    "cumulative_path_um", "z_focus", "n_planes",
]

ATOL_UM = 1e-6


def load_config(path):
    path = Path(path)
    user = json.loads(path.read_text(encoding="utf-8"))
    unknown = set(user) - set(DEFAULTS)
    if unknown:
        raise ValueError(f"Unknown config keys: {sorted(unknown)}")
    cfg = {**DEFAULTS, **user}
    # Relative paths in the config are relative to the config file.
    base = path.resolve().parent
    cfg["annotation_dirs"] = [str(p if Path(p).is_absolute() else base / p)
                              for p in cfg["annotation_dirs"]]
    if not Path(cfg["output_dir"]).is_absolute():
        cfg["output_dir"] = str(base / cfg["output_dir"])
    if cfg["index_xml_path"] and not Path(cfg["index_xml_path"]).is_absolute():
        cfg["index_xml_path"] = str(base / cfg["index_xml_path"])
    if not cfg["annotation_dirs"]:
        raise ValueError("annotation_dirs is empty")
    return cfg


# ------------------------------------------------------------------
# Operetta metadata
# ------------------------------------------------------------------

def find_index_xml(start_dir, cfg):
    if cfg["index_xml_path"]:
        return Path(cfg["index_xml_path"])
    names = {n.lower() for n in cfg["index_xml_names"]}
    current = start_dir
    for _ in range(cfg["index_search_depth"] + 1):
        for candidate in current.glob("*.xml"):
            if candidate.name.lower() in names:
                return candidate
        if current.parent == current:
            break
        current = current.parent
    raise FileNotFoundError(f"Operetta Index.xml not found above {start_dir}")


def read_channel_maps(root):
    maps = defaultdict(dict)
    for entry in root.findall(".//{*}Maps/{*}Map/{*}Entry"):
        if entry.get("ChannelID") is not None:
            maps[int(entry.get("ChannelID"))].update(
                {c.tag.split("}", 1)[-1]: c.text for c in entry})
    return maps


def read_pixel_size(xml_path):
    """ImageResolutionX/Y (metres) of the first channel that carries both."""
    maps = read_channel_maps(ET.parse(xml_path).getroot())
    for channel in sorted(maps):
        rx, ry = maps[channel].get("ImageResolutionX"), maps[channel].get("ImageResolutionY")
        if rx and ry:
            return float(rx) * 1e6, float(ry) * 1e6
    raise ValueError(f"No ImageResolutionX/Y in {xml_path}")


def calibrate_focus_metadata(table, xml_path, cfg):
    """Attach absolute focus position and acquisition time to each annotation.

    Annotated (timepoint, plane) indices are mapped to the sorted Operetta
    TimepointID/PlaneID of the brightfield channel of the source field
    (row/column/field parsed from the mask name, e.g. _r01c01f02).
    Missing or ambiguous metadata raises an error.
    """
    root = ET.parse(xml_path).getroot()
    meta = pd.DataFrame([{c.tag.split("}")[-1]: c.text for c in im}
                         for im in root.findall(".//{*}Images/{*}Image")])
    for key in ["Row", "Col", "FieldID", "ChannelID", "TimepointID", "PlaneID"]:
        meta[key] = pd.to_numeric(meta[key], errors="raise").astype(int)
    for key in ["PositionZ", "AbsPositionZ", "MeasurementTimeOffset"]:
        meta[key] = pd.to_numeric(meta[key], errors="raise")

    maps = read_channel_maps(root)
    keyword = cfg["brightfield_channel_keyword"].lower()
    bf = [c for c, m in maps.items() if any(keyword in str(v).lower() for v in m.values())]
    if len(bf) != 1:
        raise ValueError(f"Expected one '{keyword}' channel, found {bf}: {xml_path}")
    channel = bf[0]
    thr = cfg["z_jump_threshold_um"] + ATOL_UM

    pieces = []
    for source, g in table.groupby("source", sort=False):
        match = re.search(r"_r(\d+)c(\d+)f(\d+)", source)
        if not match:
            raise ValueError(f"Cannot resolve row/column/field in {source}")
        row, col, field = map(int, match.groups())
        m = meta[(meta.Row == row) & (meta.Col == col)
                 & (meta.FieldID == field) & (meta.ChannelID == channel)]
        if m.empty or m.duplicated(["TimepointID", "PlaneID"]).any():
            raise ValueError(f"Empty or ambiguous metadata for {source}")
        tids, pids = sorted(m.TimepointID.unique()), sorted(m.PlaneID.unique())
        lookup = m.set_index(["TimepointID", "PlaneID"])
        t0 = float(lookup.loc[(tids[0], pids[0]), "MeasurementTimeOffset"])

        records = []
        for _, r in g.iterrows():
            t, z = int(r.timepoint), int(r.z_focus)
            if not (0 <= t < len(tids) and 0 <= z < len(pids)):
                raise ValueError(f"Annotation outside metadata: {source}, T={t}, Z={z}")
            focus, base = lookup.loc[(tids[t], pids[z])], lookup.loc[(tids[t], pids[0])]
            rec = r.to_dict()
            rec.update(
                acquisition_time_h=(float(base.MeasurementTimeOffset) - t0) / 3600,
                focus_acquisition_time_h=(float(focus.MeasurementTimeOffset) - t0) / 3600,
                operetta_timepoint_id=int(tids[t]), operetta_plane_id=int(pids[z]),
                focus_stack_offset_um=(float(focus.PositionZ) - float(base.PositionZ)) * 1e6,
                focus_absolute_um=float(focus.AbsPositionZ) * 1e6,
                stack_origin_absolute_um=float(base.AbsPositionZ) * 1e6,
                pixel_size_x_um=float(maps[channel]["ImageResolutionX"]) * 1e6,
                pixel_size_y_um=float(maps[channel]["ImageResolutionY"]) * 1e6,
                metadata_channel_id=channel, metadata_file=Path(xml_path).name)
            records.append(rec)

        a = pd.DataFrame(records).sort_values(["label", "timepoint"])
        gb = a.groupby("label", sort=False)
        for value, prefix in [("focus_stack_offset_um", "focus_stack"),
                              ("focus_absolute_um", "focus_absolute"),
                              ("stack_origin_absolute_um", "stack_origin")]:
            a[prefix + "_step_um"] = gb[value].diff()
            a[prefix + "_change_um"] = a[value] - gb[value].transform("first")
        a["large_focus_index_jump"] = a.focus_stack_step_um.abs() > thr
        a["large_focus_absolute_jump"] = a.focus_absolute_step_um.abs() > thr
        pieces.append(a)
    return pd.concat(pieces, ignore_index=True)


# ------------------------------------------------------------------
# Masks and morphology
# ------------------------------------------------------------------

def resolve_axes(file_axes, ndim, cfg):
    """Declare axis meaning; tifffile often labels an unnamed axis "C" or "Q"."""
    override = cfg["mask_axes_override"]
    if override:
        if len(override) != ndim:
            raise ValueError(f"mask_axes_override '{override}' does not match ndim {ndim}")
        return override
    if ndim == 2:
        return "YX"
    if ndim == 3:
        leading = file_axes[0] if file_axes and file_axes[0] in "TZ" else None
        return (leading or cfg["ambiguous_axis"]) + "YX"
    if ndim == 4:
        return file_axes if file_axes and set(file_axes) >= {"T", "Z"} else "TZYX"
    raise ValueError(f"Unsupported mask ndim: {ndim}")


def load_mask(path, cfg):
    with tifffile.TiffFile(path) as tif:
        series = tif.series[0]
        data, file_axes = series.asarray(), series.axes
    axes = resolve_axes(file_axes, data.ndim, cfg)
    if data.ndim == 2:
        data, axes = data[np.newaxis, ...], "Z" + axes
    return data, axes, file_axes


def measure_plane(plane, px_x, px_y, cfg):
    if not np.any(plane != cfg["background_label"]):
        return pd.DataFrame()
    df = pd.DataFrame(regionprops_table(plane, properties=BASE_PROPERTIES))
    if cfg["min_instance_area_px"] > 0:
        df = df[df["area"] >= cfg["min_instance_area_px"]]
    if df.empty:
        return df
    px_mean = (px_x + px_y) / 2.0
    df["area_um2"] = df["area"] * px_x * px_y
    df["perimeter_um"] = df["perimeter"] * px_mean
    df["axis_major_um"] = df["axis_major_length"] * px_mean
    df["axis_minor_um"] = df["axis_minor_length"] * px_mean
    df["equivalent_diameter_um"] = df["equivalent_diameter_area"] * px_mean
    df["centroid_y_um"] = df["centroid-0"] * px_y
    df["centroid_x_um"] = df["centroid-1"] * px_x
    with np.errstate(divide="ignore", invalid="ignore"):
        df["circularity"] = np.where(df["perimeter"] > 0,
                                     4.0 * np.pi * df["area"] / np.square(df["perimeter"]), np.nan)
        df["aspect_ratio"] = np.where(df["axis_minor_length"] > 0,
                                      df["axis_major_length"] / df["axis_minor_length"], np.nan)
    df["orientation_deg"] = np.degrees(df["orientation"])
    return df


def measure_z_stack(labels_zyx, px_x, px_y, cfg):
    """One row per object, taken from the plane where it was drawn (largest area)."""
    per_plane = []
    for z in range(labels_zyx.shape[0]):
        df = measure_plane(labels_zyx[z], px_x, px_y, cfg)
        if not df.empty:
            df.insert(0, "z", z)
            per_plane.append(df)
    if not per_plane:
        return pd.DataFrame()
    stacked = pd.concat(per_plane, ignore_index=True)
    collapsed = stacked.loc[stacked.groupby("label")["area"].idxmax()].copy()
    counts = stacked.groupby("label").agg(n_planes=("z", "nunique"),
                                          z_min=("z", "min"), z_max=("z", "max"))
    return collapsed.merge(counts, on="label", how="left").rename(columns={"z": "z_focus"})


def measure_mask(labels, axes, px_x, px_y, cfg):
    rows = []
    if "T" in axes:
        src = (axes.index("T"), axes.index("Z")) if "Z" in axes else axes.index("T")
        dst = (0, 1) if "Z" in axes else 0
        moved = np.moveaxis(labels, src, dst)
        measure = measure_z_stack if "Z" in axes else measure_plane
        for t in range(moved.shape[0]):
            df = measure(moved[t], px_x, px_y, cfg)
            if not df.empty:
                df.insert(0, "timepoint", t)
                rows.append(df)
        n_timepoints = moved.shape[0]
    else:
        df = measure_z_stack(np.moveaxis(labels, axes.index("Z"), 0), px_x, px_y, cfg)
        if not df.empty:
            df.insert(0, "timepoint", 0)
            rows.append(df)
        n_timepoints = 1
    return (pd.concat(rows, ignore_index=True) if rows else pd.DataFrame()), n_timepoints


def add_xy_metrics(table, dt_h):
    """XY centroid motion in image coordinates; no drift correction implied."""
    df = table.sort_values(["label", "timepoint"]).copy()
    df["time_h"] = df["timepoint"] * dt_h
    gb = df.groupby("label", sort=False)
    df["step_um"] = np.hypot(gb["centroid_x_um"].diff(), gb["centroid_y_um"].diff())
    df["displacement_from_origin_um"] = np.hypot(
        df["centroid_x_um"] - gb["centroid_x_um"].transform("first"),
        df["centroid_y_um"] - gb["centroid_y_um"].transform("first"))
    df["cumulative_path_um"] = df.groupby("label")["step_um"].transform(
        lambda s: s.fillna(0).cumsum())
    return df


# ------------------------------------------------------------------
# QC reports (printed only; no observation is removed)
# ------------------------------------------------------------------

def report_continuity(table, n_timepoints):
    if table.empty or n_timepoints < 2:
        return
    problems = []
    for label, g in table.groupby("label"):
        present = np.sort(g["timepoint"].unique())
        missing = sorted(set(range(present[0], present[-1] + 1)) - set(present.tolist()))
        if missing:
            problems.append(f"label {int(label)}: gaps at {','.join(map(str, missing))}")
        elif present[-1] != n_timepoints - 1:
            problems.append(f"label {int(label)}: ends at t={int(present[-1])}")
    print(f"  ! {len(problems)} object(s) not continuous:" if problems
          else "  all objects continuous to the last timepoint")
    for line in problems:
        print(f"      {line}")


def report_multi_plane(table):
    if "n_planes" not in table:
        return
    multi = table[table["n_planes"] > 1]
    for _, r in multi.iterrows():
        print(f"  note: label {int(r['label'])} at t={int(r['timepoint'])} spans planes "
              f"{int(r['z_min'])}-{int(r['z_max'])}; plane {int(r['z_focus'])} used")


def report_circularity(table):
    """Pixel-based perimeters can make small, round objects exceed 1."""
    over = table[table["circularity"] > 1]
    for _, r in over.iterrows():
        print(f"  note: label {int(r['label'])} at t={int(r['timepoint'])}: "
              f"circularity {r['circularity']:.4f} > 1 (discrete perimeter estimate)")


def report_z_jumps(table, cfg):
    jumps = table[table["large_focus_index_jump"] | table["large_focus_absolute_jump"]]
    if jumps.empty:
        return
    print(f"  ! {len(jumps)} focus/stack QC event(s) above {cfg['z_jump_threshold_um']} um")
    for _, r in jumps.iterrows():
        print(f"      label {int(r['label'])} at t={int(r['timepoint'])}: "
              f"stack-relative {r['focus_stack_step_um']:+.3f} um; "
              f"stack origin {r['stack_origin_step_um']:+.3f} um; "
              f"absolute focus {r['focus_absolute_step_um']:+.3f} um")


# ------------------------------------------------------------------
# Run
# ------------------------------------------------------------------

def series_name(annotation_dir, cfg):
    if cfg["series_name"]:
        return cfg["series_name"]
    generic = {n.lower() for n in cfg["generic_folder_names"]}
    for candidate in [annotation_dir, *annotation_dir.parents]:
        if candidate.name and candidate.name.lower() not in generic:
            return candidate.name
    return annotation_dir.name


def annotation_level(mask_path, cfg):
    hit = any(re.search(p, mask_path.name, flags=re.IGNORECASE)
              for p in cfg["cluster_level_patterns"])
    return "cluster" if hit else "cell"


def finalize(table, source, xml_path, n_timepoints, cfg):
    table = add_xy_metrics(table, cfg["time_interval_h"])
    report_continuity(table, n_timepoints)
    report_multi_plane(table)
    report_circularity(table)
    table.insert(0, "source", source)
    table = table[[c for c in OUTPUT_COLUMNS if c in table]]
    result = calibrate_focus_metadata(table, xml_path, cfg)
    report_z_jumps(result, cfg)
    return result


def process_dir(annotation_dir, cfg):
    if not annotation_dir.exists():
        raise FileNotFoundError(f"Folder does not exist: {annotation_dir}")
    mask_files = sorted(annotation_dir.glob(cfg["mask_pattern"]))
    if not mask_files:
        warnings.warn(f"No files matching '{cfg['mask_pattern']}' in {annotation_dir}")
        return []

    print("=" * 70, annotation_dir, "=" * 70, sep="\n")
    xml_path = find_index_xml(annotation_dir, cfg)
    px_x, px_y = read_pixel_size(xml_path)
    print(f"Pixel size from {xml_path.name}: {px_x:.6f} x {px_y:.6f} um")
    out_dir = Path(cfg["output_dir"]) / series_name(annotation_dir, cfg)
    out_dir.mkdir(parents=True, exist_ok=True)

    results, grouped = [], []
    for mask_path in mask_files:
        labels, axes, file_axes = load_mask(mask_path, cfg)
        table, n_timepoints = measure_mask(labels, axes, px_x, px_y, cfg)
        ids = np.unique(labels)
        n_labels = int((ids != cfg["background_label"]).sum())
        level = annotation_level(mask_path, cfg)
        print(f"{mask_path.name}\n  file axes: {file_axes} -> read as: {axes}   "
              f"shape: {labels.shape}\n  labels: {n_labels}   timepoints: {n_timepoints}   "
              f"annotated as: {level}")
        if table.empty:
            warnings.warn(f"No objects found in {mask_path.name}")
            continue
        table["annotation_level"] = level
        table["n_labels_in_source"] = n_labels

        if cfg["timepoint_pattern"]:
            match = re.search(cfg["timepoint_pattern"], mask_path.name)
            if not match:
                raise ValueError(f"timepoint_pattern did not match '{mask_path.name}'")
            table["timepoint"] = int(match.group(1))
            grouped.append(table)
            continue

        result = finalize(table, mask_path.stem, xml_path, n_timepoints, cfg)
        out_path = out_dir / f"{mask_path.stem}_metrics.csv"
        if cfg["overwrite"] or not out_path.exists():
            result.to_csv(out_path, index=False)
        results.append(result)
        print()

    if cfg["timepoint_pattern"] and grouped:
        series = pd.concat(grouped, ignore_index=True)
        name = series_name(annotation_dir, cfg)
        result = finalize(series, name, xml_path, int(series["timepoint"].max()) + 1, cfg)
        result.to_csv(out_dir / f"{name}_metrics.csv", index=False)
        results.append(result)
    return results


def main():
    p = argparse.ArgumentParser(description=__doc__,
                                formatter_class=argparse.RawDescriptionHelpFormatter)
    p.add_argument("config", type=Path, help="JSON config (see config.json)")
    cfg = load_config(p.parse_args().config)

    tables = []
    for d in cfg["annotation_dirs"]:
        tables.extend(process_dir(Path(d), cfg))
    if not tables:
        print("No objects measured.")
        return

    combined = pd.concat(tables, ignore_index=True)
    out = Path(cfg["output_dir"])
    out.mkdir(parents=True, exist_ok=True)
    combined.to_csv(out / "ALL_metrics.csv", index=False)
    print("=" * 70)
    print(f"rows (timepoint x object): {len(combined)}")
    for level, g in combined.groupby("annotation_level"):
        print(f"  {level}: {g.groupby(['source', 'label']).ngroups} object(s)")
    print(f"Written to: {out}")


if __name__ == "__main__":
    main()
```
