## Supplementary_Softwares for "Spheroid surgery: a first-generation micromanipulation platform for extracting viable cell clusters from the outer layers of hydrogel-embedded spheroids": make_figure_outcomes.py

```
#!/usr/bin/env python3
"""
Outcome figure for the spheroid cell-extraction manuscript.

  a  Outcome of all extraction attempts (nested doughnut: class / code)
  b  Outcome of attempts in the final series, after parameter optimization
  c  Operating parameters (optimal vs suboptimal)

Usage
    python make_figure_outcomes.py
    python make_figure_outcomes.py --data extraction_attempts.csv --outdir figure

Input
    extraction_attempts.csv, one row per extraction attempt, with at least the
    columns `date` (YYYY-MM-DD) and `code` (T1-T4, C1-C4, S, E+, E-).

Output (in --outdir)
    Figure_outcomes.pdf / .svg / .eps   vector, editable text (TrueType / Type 42)
    Figure_outcomes.tif                 600 dpi, LZW
    Figure_outcomes_preview.png         450 dpi

Requirements
    Python >= 3.9, pandas, matplotlib >= 3.7 (Pillow is installed with matplotlib).
    Font: Arial is used when installed; otherwise Liberation Sans (metric-identical
    to Arial), then Helvetica. Layout is identical with any of these.
"""
import argparse
import math
import os
import re

import pandas as pd
import matplotlib
matplotlib.use('Agg')
import matplotlib.pyplot as plt
from matplotlib import font_manager
from matplotlib.patches import Wedge, Rectangle

# --------------------------------------------------------------------------- settings
# Start of the final experimental series: the protocol was finalized during the
# break between 3 and 12 August 2026; panel b includes every attempt from this date.
FINAL_SERIES_START = '2026-08-12'

# Panel c: operating parameters. A second line (after \n) is printed smaller.
PARAM_COLUMNS = ['Capillary tip (outer Ø)', 'Pressure', 'Release distance', 'Crosslinker volume']
PARAM_OPTIMAL = ['4–10 µm', '−200 to +300 mbar', '> ~50 µm\n(no re-attachment)', '1.35 µL']
PARAM_SUBOPTIMAL = ['> 14 µm or blunt tip', '> 500 mbar', '< 50 µm\n(risk of re-adhesion)',
                    '> 1.35 µL\n(possibly too stiff,\nhard to navigate)']

# Outcome classes and codes (order = order in the doughnut and the count table)
CLASS_ORDER = ['T', 'C', 'S', 'E']
CODES = {'T': ['T4', 'T2', 'T1', 'T3'], 'C': ['C1', 'C3', 'C2', 'C4'], 'S': ['S'], 'E': ['E+', 'E-']}
CLASS_NAME = {'T': 'Technical failure', 'C': 'Isolated, not evaluable', 'S': 'Placed, no dye',
              'E': 'DRAQ7-evaluated'}
CODE_DESC = {'T4': 'target did not detach', 'T2': 'capillary clogged', 'T1': 'capillary broke',
             'T3': 'gel too stiff', 'C1': 'lost / drift', 'C3': 'not released', 'C2': 're-adhered',
             'C4': 'damaged morphology', 'S': 'placed, no dye', 'E+': 'membrane-intact',
             'E-': 'all cells DRAQ7+'}

# Colours
BLACK = '#000000'
COL = {'T': '#eb6834', 'C': '#b3b9c0', 'S': '#2a78d6', 'E': '#1baf7a'}

# Text sizes (pt): small / body / panel title / panel letter
FS_S, FS, FS_T, FS_P = 8, 8.5, 9, 10

# Figure size (mm); all positions below are in mm
W_MM, H_MM = 180, 124


# --------------------------------------------------------------------------- helpers
def pick_font():
    available = {f.name for f in font_manager.fontManager.ttflist}
    for name in ('Arial', 'Liberation Sans', 'Helvetica'):
        if name in available:
            return name
    print('Warning: Arial, Liberation Sans and Helvetica not found; using the matplotlib default font.')
    return 'DejaVu Sans'


def tint(hexc, f):
    """Mix a colour with white (f = 0 keeps the colour, f = 1 gives white)."""
    h = hexc.lstrip('#')
    r, g, b = (int(h[i:i + 2], 16) for i in (0, 2, 4))
    r, g, b = (round(v + (255 - v) * f) for v in (r, g, b))
    return f'#{r:02x}{g:02x}{b:02x}'


E_MINUS = tint(COL['E'], 0.62)


def face(code):
    """Fill colour of an outcome code."""
    c = code[0]
    if code == 'E-':
        return E_MINUS
    if code in ('S', 'E+'):
        return COL[c]
    return tint(COL[c], [0.0, 0.25, 0.45, 0.62][CODES[c].index(code)])


def ink_on(fc):
    """Black text on light fills, white text on dark fills."""
    h = fc.lstrip('#')
    r, g, b = (int(h[i:i + 2], 16) / 255 for i in (0, 2, 4))
    return BLACK if (0.299 * r + 0.587 * g + 0.114 * b) > 0.62 else '#ffffff'


def disp(code):
    return code.replace('E-', 'E–')


def date_range_label(d0, d1):
    if d0.year == d1.year:
        return f'{d0.day} {d0:%b} – {d1.day} {d1:%b %Y}'
    return f'{d0.day} {d0:%b %Y} – {d1.day} {d1:%b %Y}'


# --------------------------------------------------------------------------- figure
def make_figure(df, outdir):
    font = pick_font()
    plt.rcParams.update({'font.family': font, 'font.size': FS, 'svg.fonttype': 'none',
                         'pdf.fonttype': 42, 'ps.fonttype': 42, 'axes.linewidth': 0.5,
                         'lines.linewidth': 0.5, 'text.color': BLACK})

    N = len(df)
    cc = df.cls.value_counts()
    cd = df.code.value_counts()

    fig = plt.figure(figsize=(W_MM / 25.4, H_MM / 25.4))

    def axmm(x, y, w, h):
        a = fig.add_axes([x / W_MM, y / H_MM, w / W_MM, h / H_MM])
        a.axis('off')
        return a

    def panel(x, y, s, title, subtitle=None):
        fig.text(x / W_MM, y / H_MM, s, fontsize=FS_P, fontweight='bold', va='top', ha='left')
        fig.text((x + 4.2) / W_MM, (y - 0.35) / H_MM, title, fontsize=FS_T, fontweight='bold', va='top', ha='left')
        if subtitle:
            fig.text((x + 4.2) / W_MM, (y - 4.4) / H_MM, subtitle, fontsize=FS_S, va='top', ha='left')

    # ----------------------------------------------------------------- (a) doughnut
    panel(0, 123, 'a', f'Outcome of all extraction attempts (n = {N})')
    ax = axmm(0, 54, 61, 61)
    ax.set_aspect('equal')
    ax.set_xlim(-1.03, 1.03)
    ax.set_ylim(-1.03, 1.03)
    R_in0, R_in1, R_out0, R_out1 = 0.40, 0.68, 0.71, 1.0

    def ang(frac):  # clockwise from 12 o'clock
        return 90 - 360 * frac

    acc = 0
    for c in CLASS_ORDER:  # inner ring: classes
        n = int(cc.get(c, 0))
        if n == 0:
            continue
        a0, a1 = ang((acc + n) / N), ang(acc / N)
        ax.add_patch(Wedge((0, 0), R_in1, a0, a1, width=R_in1 - R_in0, facecolor=COL[c],
                           edgecolor='white', linewidth=0.8))
        m = math.radians((a0 + a1) / 2)
        rr = (R_in0 + R_in1) / 2
        ax.text(rr * math.cos(m), rr * math.sin(m), f'{c}\n{n}', ha='center', va='center', fontsize=FS,
                fontweight='bold', color=ink_on(COL[c]), linespacing=1.0)
        acc += n

    acc = 0
    for c in CLASS_ORDER:  # outer ring: codes
        for d in CODES[c]:
            n = int(cd.get(d, 0))
            if n == 0:
                continue
            a0, a1 = ang((acc + n) / N), ang(acc / N)
            fc = face(d)
            ax.add_patch(Wedge((0, 0), R_out1, a0, a1, width=R_out1 - R_out0, facecolor=fc,
                               edgecolor='white', linewidth=0.5))
            m = math.radians((a0 + a1) / 2)
            rr = (R_out0 + R_out1) / 2
            ax.text(rr * math.cos(m), rr * math.sin(m), f'{n}', ha='center', va='center',
                    fontsize=FS_S if n < 2 else FS, fontweight='bold', color=ink_on(fc))
            acc += n
    ax.text(0, 0.09, f'{N}', ha='center', va='center', fontsize=FS_T, fontweight='bold')
    ax.text(0, -0.12, 'attempts', ha='center', va='center', fontsize=FS_S)

    # count table next to the doughnut
    tb = axmm(62, 54, 54, 60)
    tb.set_xlim(0, 54)
    tb.set_ylim(0, 60)
    y = 58.4
    rowh = 4.0
    for c in CLASS_ORDER:
        tb.add_patch(Rectangle((0, y - 1.3), 2.6, 2.6, facecolor=COL[c], edgecolor='none'))
        tb.text(3.8, y, c, va='center', fontsize=FS, fontweight='bold')
        tb.text(7.6, y, CLASS_NAME[c], va='center', fontsize=FS, fontweight='bold')
        tb.text(54, y, f'{int(cc.get(c, 0))}', va='center', ha='right', fontsize=FS, fontweight='bold')
        y -= rowh
        if c == 'S':
            y -= 0.8
            continue
        for d in CODES[c]:
            tb.add_patch(Rectangle((3.9, y - 1.1), 2.2, 2.2, facecolor=face(d), edgecolor='none'))
            tb.text(7.6, y, disp(d), va='center', fontsize=FS, fontweight='bold')
            tb.text(13.0, y, CODE_DESC[d], va='center', fontsize=FS_S)
            tb.text(54, y, f'{int(cd.get(d, 0))}', va='center', ha='right', fontsize=FS)
            y -= rowh
        y -= 0.8

    # ----------------------------------------------------------------- (b) final series
    start = pd.Timestamp(FINAL_SERIES_START)
    p = df[df.date >= start]
    n0 = len(p)
    k = {'T': int((p.cls == 'T').sum()), 'C': int((p.cls == 'C').sum()), 'S': int((p.cls == 'S').sum()),
         'E+': int((p.code == 'E+').sum()), 'E-': int((p.code == 'E-').sum())}
    panel(120, 123, 'b', 'After parameter optimization',
          f'All attempts, {date_range_label(p.date.min(), p.date.max())} · n = {n0}')
    ax = axmm(119, 54, 61, 58)
    ax.set_xlim(0, 61)
    ax.set_ylim(0, 58)
    y0, y1 = 2, 54
    u = (y1 - y0) / n0
    xa, x0, x1 = 10, 14, 29
    ax.plot([xa, xa], [y0, y1], color=BLACK, lw=0.5)
    for t in range(0, n0 + 1, 5):
        yy = y0 + t * u
        ax.plot([xa - 1.0, xa], [yy, yy], color=BLACK, lw=0.5)
        ax.text(xa - 1.7, yy, f'{t}', ha='right', va='center', fontsize=FS_S)
    ax.text(1.3, (y0 + y1) / 2, 'Attempts (n)', rotation=90, ha='center', va='center', fontsize=FS)

    segs = {}
    y = y0
    for c in ['E+', 'E-', 'S', 'C', 'T']:  # bottom to top
        if k[c] == 0:
            continue
        h = k[c] * u
        segs[c] = (y, y + h)
        fc = face(c) if c not in ('C', 'T') else COL[c]
        ax.add_patch(Rectangle((x0, y), x1 - x0, h, facecolor=fc, edgecolor='white', linewidth=0.6))
        ax.text((x0 + x1) / 2, y + h / 2, f'{disp(c)}  {k[c]}', ha='center', va='center', fontsize=FS,
                fontweight='bold', color=ink_on(fc))
        y += h

    def bracket(ya, yb, pct, line1, line2=None):
        xb = x1 + 1.6
        ax.plot([xb, xb + 1.0, xb + 1.0, xb], [ya + 0.35, ya + 0.35, yb - 0.35, yb - 0.35], color=BLACK, lw=0.5)
        ym = (ya + yb) / 2
        tx = xb + 2.4
        ax.text(tx, ym + (3.0 if line2 else 1.6), pct, ha='left', va='center', fontsize=FS, fontweight='bold')
        ax.text(tx, ym - (0.2 if line2 else 1.6), line1, ha='left', va='center', fontsize=FS_S)
        if line2:
            ax.text(tx, ym - 3.4, line2, ha='left', va='center', fontsize=FS_S)

    placed = k['E+'] + k['E-'] + k['S']
    placed_keys = [c for c in ['E+', 'E-', 'S'] if k[c]]
    if placed:
        bracket(segs[placed_keys[0]][0], segs[placed_keys[-1]][1], f'{placed / n0 * 100:.0f}% ({placed}/{n0})',
                'placed in gel', f'E+ {k["E+"]}/{k["E+"] + k["E-"]} evaluated' if k['E+'] + k['E-'] else None)
    if k['C']:
        bracket(*segs['C'], f'{k["C"] / n0 * 100:.0f}% ({k["C"]}/{n0})', 'not evaluable')
    if k['T']:
        bracket(*segs['T'], f'{k["T"] / n0 * 100:.0f}% ({k["T"]}/{n0})', 'technical failure')

    # ----------------------------------------------------------------- (c) parameters
    panel(0, 49, 'c', 'Operating parameters')
    T = axmm(0, 1, 180, 41)
    T.set_xlim(0, 180)
    T.set_ylim(0, 41)
    rowlab = 21
    colw = (180 - rowlab) / len(PARAM_COLUMNS)
    top, hdr, rowh = 40.5, 6.5, 15.5
    T.plot([0, 180], [top, top], color=BLACK, lw=0.6)
    for j, c in enumerate(PARAM_COLUMNS):
        T.text(rowlab + j * colw + 1.5, top - hdr / 2, c, fontsize=FS, fontweight='bold', va='center', ha='left')
    T.plot([0, 180], [top - hdr, top - hdr], color=BLACK, lw=0.5)
    for i, (rl, col, vals) in enumerate([('Optimal', COL['E'], PARAM_OPTIMAL),
                                         ('Suboptimal', COL['T'], PARAM_SUBOPTIMAL)]):
        yt = top - hdr - i * rowh
        yb = yt - rowh
        ym = (yt + yb) / 2
        T.add_patch(Rectangle((0, yb + 0.4), 180, rowh - 0.8, facecolor=tint(col, 0.88), edgecolor='none', zorder=0))
        T.add_patch(Rectangle((0, yb + 0.4), 1.3, rowh - 0.8, facecolor=col, edgecolor='none', zorder=1))
        T.text(3.2, ym, rl, fontsize=FS, fontweight='bold', va='center', ha='left')
        for j, v in enumerate(vals):
            main, *rest = v.split('\n')
            x = rowlab + j * colw + 1.5
            if rest:
                top_y = ym + (1.9 + 1.7 * (len(rest) - 1))
                T.text(x, top_y, main, fontsize=FS, fontweight='bold', va='center', ha='left')
                T.text(x, top_y - 2.0, '\n'.join(rest), fontsize=FS_S, va='top', ha='left', linespacing=1.15)
            else:
                T.text(x, ym, main, fontsize=FS, fontweight='bold', va='center', ha='left')
    T.plot([0, 180], [top - hdr - 2 * rowh, top - hdr - 2 * rowh], color=BLACK, lw=0.6)

    # ----------------------------------------------------------------- save
    os.makedirs(outdir, exist_ok=True)
    base = os.path.join(outdir, 'Figure_outcomes')
    kw = dict(bbox_inches='tight', pad_inches=0.04)
    fig.savefig(base + '.pdf', metadata={'CreationDate': None}, **kw)
    fig.savefig(base + '.svg', metadata={'Date': None}, **kw)
    fig.savefig(base + '.eps', **kw)
    fig.savefig(base + '.tif', dpi=600, pil_kwargs={'compression': 'tiff_lzw'}, **kw)
    fig.savefig(base + '_preview.png', dpi=450, **kw)
    plt.close(fig)

    # Name the font Arial in the SVG so it opens as Arial in Illustrator / Inkscape
    with open(base + '.svg', encoding='utf-8') as f:
        svg = f.read()
    svg = svg.replace(f"'{font}'", 'Arial').replace(font, 'Arial')
    with open(base + '.svg', 'w', encoding='utf-8') as f:
        f.write(svg)

    sizes = sorted(set(float(x) for x in re.findall(r'font-size: ([0-9.]+)px', svg)))
    print(f'Font: {font}; text sizes (pt): {sizes}')
    print(f'Panel a: n = {N}; classes { {c: int(v) for c, v in cc.items()} }')
    print(f'Panel b: n = {n0} from {FINAL_SERIES_START}; {k}')
    print(f'Saved: {base}.pdf / .svg / .eps / .tif / _preview.png')


def main():
    here = os.path.dirname(os.path.abspath(__file__))
    ap = argparse.ArgumentParser(description=__doc__, formatter_class=argparse.RawDescriptionHelpFormatter)
    ap.add_argument('--data', default=os.path.join(here, 'extraction_attempts.csv'), help='attempt-level CSV')
    ap.add_argument('--outdir', default=os.path.join(here, 'figure'), help='output folder')
    args = ap.parse_args()

    df = pd.read_csv(args.data, dtype={'code': str})
    df['code'] = df['code'].str.strip().str.replace('–', '-', regex=False)
    df['date'] = pd.to_datetime(df['date'])
    known = {c for v in CODES.values() for c in v}
    unknown = sorted(set(df.code) - known)
    if unknown:
        raise SystemExit(f'Unknown outcome codes in {args.data}: {unknown}')
    df['cls'] = df.code.str[0]
    make_figure(df, args.outdir)


if __name__ == '__main__':
    main()
```
