## Supplementary material for "Spheroid surgery: a first-generation micromanipulation platform for extracting viable cell clusters from the outer layers of hydrogel-embedded spheroids": Supplemetary_Note

### Supplementary Note 1

One block per sample (header row: cell line and crosslinker volume), one row per extraction attempt in chronological order. Each attempt is assigned exactly one outcome code. Each sample identifier corresponds to a distinct spheroid-containing hydrogel preparation.

#### 1. Outcome classification

| Class | Code | Definition |
|---|---|---|
| T | T1 | Capillary broke |
| | T2 | Capillary clogged |
| | T3 | Gel not navigable (too stiff) |
| | T4 | Target did not detach from spheroid |
| C | C1 | Lost from field of view / drift |
| | C2 | Re-adhered to spheroid or other cells |
| | C3 | Cannot release the cell from the pipette |
| | C4 | Cell looked damaged by morphology |
| E | E+ | DRAQ7-negative for at least one cell throughout the observation window (membrane-intact) |
| | E– | All identifiable cells within the extracted unit became DRAQ7-positive during the observation window; the time of detection was recorded. |
| S | S | Successful extraction without screening with dye |

T = technical failure (no isolated unit obtained). C = extracted but not evaluable (censored). E = evaluable.

#### 2. Summary

| Specimens | Attempts | T total | C total | E total | S total | T1 | T2 | T3 | T4 | C1 | C2 | C3 | C4 | E+ | E– | S |
|---|---|---|---|---|---|---|---|---|---|---|---|---|---|---|---|---|
| 19 | 55 | 26 | 11 | 8 | 10 | 3 | 10 | 2 | 11 | 5 | 1 | 4 | 1 | 6 | 2 | 10 |

#### 3. Attempt-level log

| Attempt | Code | Video file | Type[1] | Targeted ~n cells | Dist. from spheroid (µm) | Capillary tip (µm) | Notes |
|---|---|---|---|---|---|---|---|
| **20260723_H_01** | HeLa Kyoto · crosslinker 1.50 µL | | | | | | |
| 01 | S | 260723155121-H-S-01.mp4 | singlet | 3 | ~70 | ~4.5 | Tip measured on a slightly blurred frame |
| 02 | S | 260723161203-H-S-02.mp4 | singlet | 1 | ~80 | ~5 | |

| | | | | | | | |
|---|---|---|---|---|---|---|---|
| **20260724_H_01** | HeLa Kyoto · crosslinker 1.40 µL | | | | | | |
| 01 | **S** | 260724145306-H-M-01.mp4 | cluster | ~3 | >300 | n.m. | Spheroid and cluster never in the same high-mag frame; tip only seen against the spheroid or covered by cells |
| **20260730_H_01** | HeLa Kyoto · crosslinker 1.35 µL | | | | | | |
| 01 | **C4** | 20260730_H_01.mp4 | cluster | 2–3 | – | ~8.5 | the cells looked damaged; blunt tip |
| 02 | **T4** | 260730140838-H-02.mp4 | – | – | – | ~5 | |
| 03 | **C3** | 260730142937-H-M-03.mp4 | cluster | ~3 | – | ~6.5 | Dark video, tip value approximate |
| 04 | **T2** | 260730143956-H-04.mp4 | – | – | – | ~5 | |
| **20260730_H_02** | HeLa Kyoto · crosslinker 1.35 µL | | | | | | |
| 01 | **T2** | 260730150121-H-05.mp4 | – | – | – | ~5.5 | |
| 02 | **C1** | 260730151457-H-M-06.mp4 | cluster | ~2 | – | ~5 | Last seen on the pipette ~75 µm from the spheroid |
| 03 | **T2** | 260730152343-H-07.mp4 | – | – | – | ~4 | |
| 04 | **T2** | 260730152811-H-08.mp4 | – | – | – | ~4.5 | |
| **20260730_H_03** | HeLa Kyoto · crosslinker 1.35 µL | | | | | | |
| 01 | **C3** | 260730153540-H-M-09.mp4 | cluster | ~6 | – | ~4.5 | |
| 02 | **C1** | 260730154627-H-M-10.mp4 | cluster | ~6 | – | ~4.5 | |
| 03 | **C3** | 260730155938-H-M-11.mp4 | cluster | ~4 | – | ~4 | |
| 04 | **T2** | 260730161118-H-12.mp4 | – | – | – | ~5 | |
| **20260731_H_01** | HeLa Kyoto · crosslinker 1.35 µL | | | | | | |
| 01 | **T2** | 260731091004-H-01.mp4 | – | – | – | ~10 | Blunt tip |
| 02 | **T2** | 260731091742-H-02.mp4 | – | – | – | ~10 | Blunt tip |
| 03 | **T4** | 260731092118-H-03.mp4 | – | – | – | ~5.5 | |
| 04 | **C1** | 260731092957-H-M-04.mp4 | cluster | ~2 | – | ~6.5 | Last seen on the pipette ~65 µm from the spheroid |
| 05 | **T3** | 260731094621-H-05.mp4 | – | – | – | ~5 | |
| **20260731_H_02** | HeLa Kyoto · crosslinker 1.35 µL | | | | | | |
| 01 | **T4** | 260731095226-H-06.mp4 | – | – | – | ~5.5 | |
| 02 | **T2** | 260731101525-H-07.mp4 | – | – | – | ~5.5 | |
| 03 | **T4** | 260731102633-H-08.mp4 | – | – | – | ~5 | |
| 04 | **T4** | 260731103650-H-09.mp4 | – | – | – | ~4.5 | |
| **20260731_H_03** | HeLa Kyoto · crosslinker 1.35 µL | | | | | | |
| 01 | **S** | 260731104751-H-M-10.mp4 | cluster | ? | >200 | ~4.5 | Target unclear: large fragment (~80 µm) or small unit on the tip. |
| 02 | **T4** | 260731114714-H-11.mp4 | – | – | – | ~4 | |
| 03 | **C2** | 260731115750-H-S-12.mp4 | singlet | 1 | 0 | ~4.5 | Re-adhered |
| 04 | **S** | 260731122808-H-S-13.mp4 | singlet | 1 | ~50 | ~4.5 | |
| **20260731_H_04** | HeLa Kyoto · crosslinker 1.35 µL | | | | | | |
| 01 | **T4** | 260731163500-H-13.mp4 | – | – | – | ~4.5 | Dark frame, tip value uncertain |

| | | | | | | | |
|---|---|---|---|---|---|---|---|
| 02 | T4 | 260731165148-H-14.mp4 | – | – | – | ~3.5 | |
| 03 | T2 | 260731165745-H-15.mp4 | – | – | – | ~4.5 | |
| 04 | S | 260731170441-H-M-16.mp4 | cluster | 2 | ~50 | ~4.5 | Spheroid extends below the frame; the gap may be smaller |
| **20260801_H_01** | | HeLa Kyoto · crosslinker 1.35 µL | | | | | |
| 01 | S | 260801150655-H-M-01.mp4 | cluster | ~3 | ~400 | ~4 | |
| 02 | T2 | 260801152454-H-02.mp4 | – | – | – | ~6 | |
| 03 | S | 260801153817-H-M-03.mp4 | cluster | 2 | ~100 | ~5.5 | Doublet in a different focal plane, overlaps the spheroid in xy |
| **20260803_H_01** | | HeLa Kyoto · crosslinker 1.35 µL | | | | | |
| 01 | C1 | 260803084508-H-M-01.mp4 | cluster | 2–3 | – | ~4.5 | |
| 02 | S | 260803092640-H-M-01.mp4 | cluster | 2 | ~350 | ~5 | |
| 03 | C3 | 260803095740-H-M-03.mp4 | cluster | ~2 | – | ~5.5 | |
| 04 | C1 | 260803100643-H-M-04.mp4 | cluster | ~3 | – | ~5 | |
| **20260803_H_02** | | HeLa Kyoto · crosslinker 1.35 µL | | | | | |
| 01 | T4 | 260803113625-H-01.mp4 | – | – | – | ~5 | |
| 02 | E+ | 20260803_H_M_02_02.mp4 | cluster | ~14 | ~200 | ~4.5 | Spheroid edge out of focus. |
| **20260803_H_03** | | HeLa Kyoto · crosslinker 1.35 µL | | | | | |
| 01 | T3 | 260803121217-H-06.mp4 | – | – | – | ~5 | |
| 02 | T4 | 260803121755-H-07.mp4 | – | – | – | ~4 | |
| 03 | T4 | 260803122502-H-08.mp4 | – | – | – | ~4.5 | |
| **20260812_H_01** | | HeLa Kyoto · crosslinker 1.35 µL | | | | | |
| 01 | E+ | 20260812_H_M_01_01.mp4 | cluster | ~3 | ~80 | ~4.5 | Elongated cell, probably 21 in the extraction 1 stayed DRAQ7 negative |
| **20260817_H_01** | | HeLa Kyoto · crosslinker 1.35 µL | | | | | |
| 01 | T1 | 260817095602-H-01.mp4 | – | – | – | ~9 | Blunt tip at the start of the video |
| 02 | T1 | 260817100905-H-02.mp4 | – | – | – | ~7 | |
| 03 | T1 | 260817110516-H-03.mp4 | – | – | – | ~4 | |
| 04 | E+ | 20260817_H_M_01_04.mp4 | cluster | ~2 | ~600 | ~4 | Cells strung along the pipette; final unit out of focus. |
| **20260901_T_01** | | T-47D · crosslinker 1.35 µL | | | | | |
| 01 | E+ | 20260901_T_D_01_01.mp4 | doublet | 4 | >50 | ~6.5 | A second small cell sits at the spheroid edge, between the target and the spheroid. |
| **20260904_T_01** | | T-47D · crosslinker 1.35 µL | | | | | |
| 01 | S | 260904101151-T-M-01.mp4 | cluster | 1–3 | ~500 | ~7.5 | Compact ~40 × 35 µm unit, cell boundaries not visible. |
| 02 | E+ | 20260904_T_D_01_02.mp4 | doublet | 3 | ~700 | ~8 | Elongated doublet. |
| **20260917_T_01** | | T-47D · crosslinker 1.35 µL | | | | | |
| 01 | E– | 20260917_T_S_01_01.mp4 | singlet | 1 | ~190 | ~14 | At the spheroid's widest plane the cell lies over the spheroid outline. |
| 02 | E– | 20260917_T_S_01_02.mp4 | singlet | 1 | ~150 | ~5 | Only one cell visible after release. |

| | | | | | | |
|---|---|---|---|---|---|---|
| **20260925_H_01** | HeLa Kyoto · crosslinker 1.35 µL | | | | | |
| 01 | **E+** | 20260925_H_M_01_01.mp4 | cluster | 3 | ~350 | ~4 |

[1] Target type for T, C and S attempts; extracted type for E attempts.

### 4. DRAQ7-evaluated extractions (E)

| Sample ID | Attempt | Video file | Code | Extracted type | Time to E− classification (h) | Window (h) | Division (first) |
|---|---|---|---|---|---|---|---|
| **20260803_H_02** | 02 | 20260803_H_M_02_02.mp4 | **E+** | cluster | – | 24 | 6 h |
| **20260812_H_01** | 01 | 20260812_H_M_01_01.mp4 | **E+** | cluster | – | 24 | – |
| **20260817_H_01** | 04 | 20260817_H_M_01_04.mp4 | **E+** | cluster | – | 24 | – |
| **20260901_T_01** | 01 | 20260901_T_D_01_01.mp4 | **E+** | doublet | – | 24 | – |
| **20260904_T_01** | 02 | 20260904_T_D_01_02.mp4 | **E+** | doublet | – | 24 | – |
| **20260917_T_01** | 01 | 20260917_T_S_01_01.mp4 | **E–** | singlet | 0 | 23 | – |
| **20260917_T_01** | 02 | 20260917_T_S_01_02.mp4 | **E–** | singlet | 0 | 23 | – |
| **20260925_H_01** | 01 | 20260925_H_M_01_01.mp4 | **E+** | cluster | – | 24 | – |

### 5. Notes

Attempt = the running number within the sample. The original day-level numbering is preserved in the video file name. In T rows the cells that describe the extracted unit are marked with a dash: the attempt failed before an isolated unit existed.

**Measurement notes.** Pixel size 0.22374 µm/px (LNscope, 640 × 480 video frames). Capillary tip = outer diameter at the tip, measured at the start of each video where the pipette is against clear gel (about ±1 µm). Dist. from spheroid = approximate shortest distance between the edge of the placed unit and the spheroid outline. Because units were displaced diagonally through the gel, this distance includes an axial component, estimated from the focus change between the placed unit and the nearest spheroid edge. Gaps within one field of view (≤ ~140 µm) were measured on the video frame; larger gaps, and all axial components, were estimated visually while moving the stage and focus, except for 20260803_H_02/02 (~200 µm), which was measured from the stage position. All distances are therefore approximate. n.m. = not measured. A dash in C rows = not applicable (unit not released or lost).
